# The INO80 ATP-dependent chromatin remodelling complex alleviates stalled Polymerase II to promote non-coding RNA transcription termination

**DOI:** 10.1101/2020.03.02.973685

**Authors:** Sara Luzzi, Ugo Szachnowski, Sarah Greener, Kenny Schumacher, Stuart Fulton, Chloe Walton, Camille Gautier, Kang Hoo Han, Jack Darke, Rossana Piccinno, Anne Lafon, B. Franklin Pugh, Didier Devys, Laszlo Tora, Antonin Morillon, Manolis Papamichos-Chronakis

## Abstract

Co-transcriptional RNA quality control is essential for gene expression. However, its regulation remains poorly understood. Here, we report that the evolutionarily conserved ATP-dependent chromatin remodelling INO80 complex promotes transcription termination by the non-coding RNA quality control pathway in *S. cerevisiae*. Loss of INO80 leads to accumulation of stalled RNA Polymerase II preferentially at promoter-proximal pausing sites, compromising Pol II processivity and hindering transcription elongation. We reveal that binding of RNA surveillance and non-coding transcription termination factors to promoter-proximal mRNA regions is associated with increased promoter-proximal pausing. INO80 counteracts promoter-proximal stalling of genes attenuated by the Nrd1-Nab3-Sen1 (NNS) non-coding transcription termination complex, promoting their expression. We show that INO80 interacts with Nrd1 and the Nab2 RNA surveillance factor *in vivo*. Absence of INO80 leads to defective transcription termination by the Nrd1-Nab3-Sen1 (NNS) complex. We demonstrate that INO80 facilitates the recruitment of Nab2 at non-coding transcription termination sites and its association with promoter-proximally terminated mRNA transcripts. Finally, we provide evidence that INO80 promotes the release of stalled RNA Polymerase II from a non-coding transcription termination site. Collectively, our work suggests that the INO80 complex regulates transcription by removal of stalled Polymerase, implicating a chromatin-based mechanism for non-coding and premature transcription termination in gene expression.

## INTRODUCTION

Elimination of aberrant mRNAs is paramount to proper gene expression. In eukaryotic cells nuclear RNA surveillance and quality control mechanisms monitor mRNA biogenesis co-transcriptionally and prematurely terminate transcription for degradation of nascent RNA transcripts^1^. Elegant genome-wide studies in the budding yeast *Saccharomyces cerevisiae* have suggested that nonproductive transcriptional elongation of protein-coding genes is wide-spread and common^2,3^. Furthermore, the identification of thousands of promoter-proximal short mRNAs enriched in 3’-oligo(A) tails^4^, indicates extensive premature termination across the yeast protein-coding genome^5^. Likewise, in human cells only 10% of promoter-proximally paused polymerases was shown to enter productive elongation, with the remaining being prematurely terminated during abortive elongation^6,7^.

A major nuclear RNA quality control pathway in yeast is mediated by the complex Nrd1-Nab3-Sen1 (NNS)^8^. NNS is aided by the nuclear RNA surveillance factor Nab2^9^ and the Trf4/Trf5-Air1/Air2-Mtr4 polyadenylation (TRAMP) complex, which participate in processing and delivering of nascent transcripts to the Rrp6-dependent nuclear exosome for degradation by its 3’ exonuclease activity^10–15^. The NNS-dependent RNA quality control pathway is essential for the termination and maturation of the structural small nucleolar RNAs (snoRNAs), while it also acts to restrict the extensive and pervasive non-coding RNA (ncRNA) transcription across the genome^16–18^. The NNS complex has been reported to prematurely terminate transcription of mRNA genes^19–21^, although a recent study suggested that transcription attenuation by NNS is rare^22^. Notably, this model has been challenged by other studies showing that several components of the non-coding termination machinery including Nrd1, Nab3, Nab2 and Mtr4 bind most mRNA transcripts and are recruited genome-wide at active protein-coding genes with a preference near their 5’ end^23–27^. These results suggest that non-coding transcription termination may play a genome-wide, yet unclear, role in regulation of mRNA gene expression^27^. Thus, although the non-coding RNA surveillance and termination pathways have been characterized in molecular detail, the mechanisms regulating premature transcription termination during mRNA synthesis remain poorly understood.

In metazoans, expression of non-coding RNAs and short promoter-associated polyadenylated mRNA transcripts is extensive and dependent on the Integrator endonuclease^28^ and the nuclear exosome^29^. Interestingly, prematurely terminated transcripts have been found to coincide with promoter-proximal Pol II pausing^30^ and experiments with chemical transcriptional inhibitors indicated high rates of turnover and premature termination of promoter-proximal paused Pol II^6,31,32^. Still, how pausing is linked to premature termination is largely unclear.

Recent evidence has indicated that premature termination of mRNA genes and promoter-proximal transcriptional pausing have been associated with nucleosome organization^30^. The nucleosomal landscape of the eukaryotic genome is shaped by ATP-dependent chromatin remodelling enzymes. Chromatin remodelers act at almost all stages of RNA production controlling transcription activation and silencing, modulation of transcriptional rates, transcription termination and RNA export from the nucleus^33–37^. However, their role in co-transcriptional RNA quality control remains elusive.

INO80 is a multisubunit ATP-dependent chromatin remodelling complex with a well-established and important role in transcription across eukaryotes^38^. INO80 is enriched at the transcription start sites (TSS) of most active and poised mRNA genes in yeast and mammals^39,40^. The nucleosome remodelling activity of INO80 is important for the exchange of the histone variant H2A.Z for H2A in nucleosomes and for controlling nucleosome positions^38^. While it is generally proposed that the function of INO80 during transcription is to control transcription initiation, INO80 has been found to be recruited not only at the TSS but also further downstream into the gene bodies^40,41^. Additionally, reports have indicated physical interaction of INO80 with the elongating RNA Polymerase II machinery and mRNA transcripts^42–45^. These observations indicate a -yet-undefined role for INO80 in the transcriptional process beyond transcription initiation.

Here we investigate the role of INO80 in transcription regulation in *S. cerevisiae*. We demonstrate that loss of INO80 leads to a global reduction in transcription and defective non-coding transcription termination. We show that INO80 alleviates stalled Pol II at DNA sequence-determined pausing sites that are defined by the ATG motif and enables the progression of Pol II from the promoter-proximal region into the gene body. Our analysis indicates that the role of INO80 in alleviating stalled Pol II is linked to loading of non-coding transcription termination and RNA processing factors to mRNAs and is associated with non-coding RNA transcription termination. Our study reveals that INO80 facilitates the release of stalled Pol II from non-coding RNA termination sites. Overall, our results suggest that the chromatin remodelling function of INO80 promotes non-coding transcription termination, coupling transcriptional pausing to RNA surveillance.

## RESULTS

### Global downregulation of Pol II transcription in the absence of INO80 is associated with defective mRNA degradation

Gene expression studies have suggested both positive and negative role for INO80 in transcription by RNA Polymerase II^38^. To shed light on the function of INO80 in Pol II transcription, we performed 4-thiouracil RNA coupled with sequencing (4tU-seq) analysis in budding yeast^46^ to evaluate RNA synthesis rates in the absence of INO80. Wild-type (WT) and *ino80*Δ cells were pulse-labeled with 4tU to label newly synthesized RNAs, and nascent and total RNA samples were collected from both strains. In addition, we performed 4tU-seq in *ino80*Δ cells expressing wild-type INO80 from a plasmid to control for *ino80*Δ-mediated changes in transcription.

By normalizing the data to the *S. pombe* spiked-in RNA signal and applying a statistical cutoff of 1.5-fold-change and p-value < 0.05, our analysis of protein-coding genes revealed that the absence of INO80 led to a significant increase in only 3.6% of the nascent transcriptome (n=178) while 34% of genes showed a significant reduction in mRNA synthesis in *ino80*Δ compared to WT (n=1706), (Fig. 1a).

**Fig. 1.**
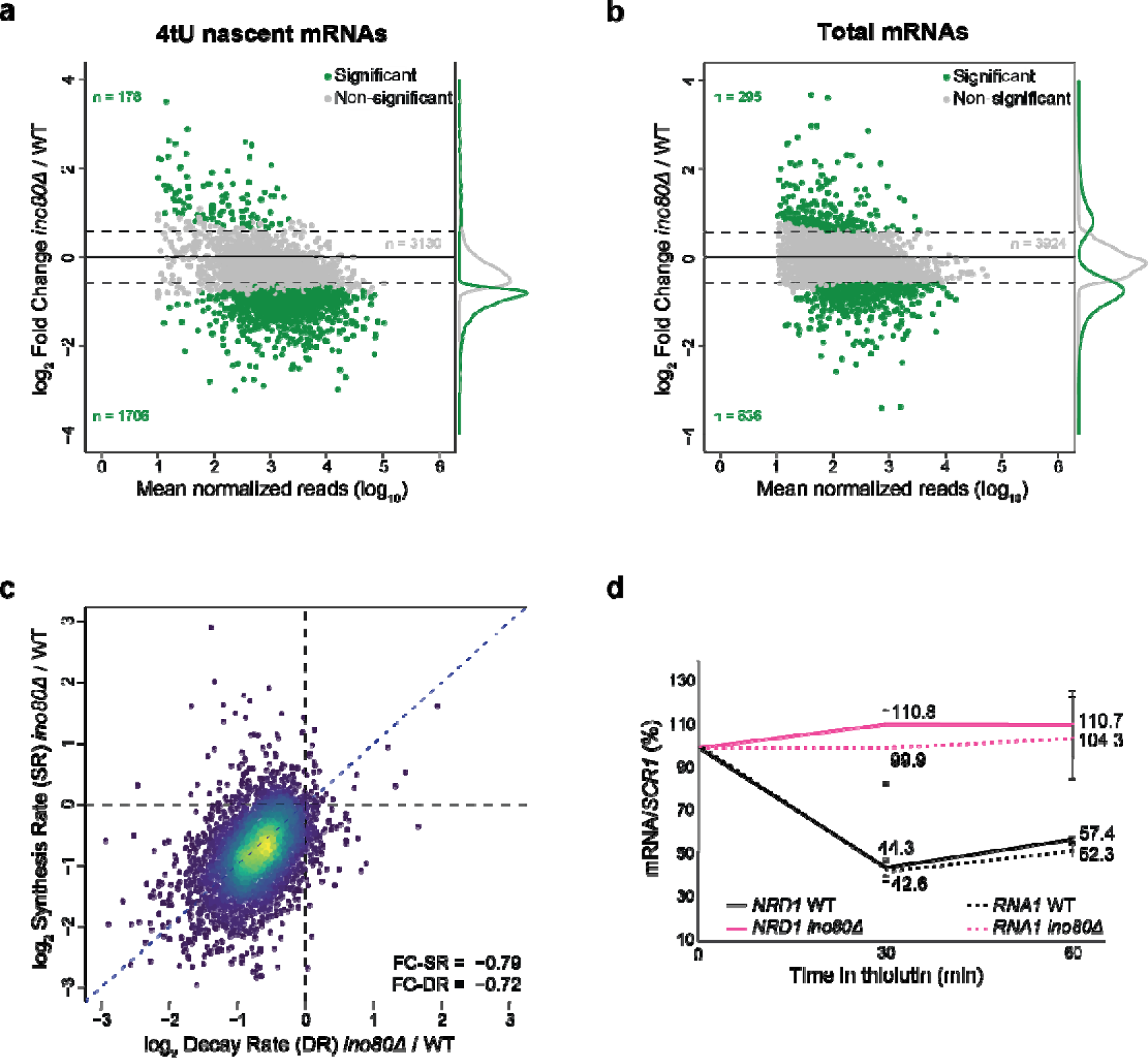
Reduced mRNA synthesis and decay rates in the absence of INO80. **a, b,** Scatterplot analysis for fold changes between WT and *ino80*Δ in newly synthesized mRNA (a) and steady-state mRNA (b) levels (n=5012 genes) plotted on a log_2_ scale. Thresholds of 1.5-fold change and 0.05 p values were considered for significant change (green dots). n=3. **c,** cDTA profiles for *ino80*Δ. For all analysed genes, changes in synthesis rates (SR) were plotted against the changes in mRNA decay rates (DR). Fold-change in SR and DR were calculated between *ino80*Δ and WT and plotted on a log_2_ scale. 90% of genes are contained within the outer contour. Yellow and red dots correspond to 60% of genes. **d,** RT-qPCR analysis for *NRD1* and *RNA1* RNAs was conducted in WT and *ino80*Δ before (T_0_) and after addition of thiolutin in YPD. RNA abundance at each time-point was normalized over the respective abundance of the *SCR1* RNA. RNA before thiolutin treatment was set at 100% in both strains. Remaining RNA was calculated as the amount of normalized RNA in the indicated time-points relative to the normalized RNA at T_0_. Values represent the average from at least three independent experiments. Bars, standard errors.

Importantly, the decrease in mRNA synthesis rates was found to be dependent on INO80, as expression of INO80 from a plasmid rescued the defect in *ino80*Δ cells, with less than 4% of genes showing decreased mRNA synthesis (n=193) (Extended Data Fig. 1a). These results indicate that deletion of INO80 globally downregulates mRNA synthesis, suggesting that INO80 plays a positive regulatory role in transcription.

Interestingly, analysis of total RNA-seq data showed that the steady-state mRNA expression levels did not significantly change for 81% of protein-coding genes in *ino80*Δ (Fig. 1b). The total RNA levels were increased for 6% of genes (n=295), while only 13% showed a significant decrease in their RNA expression levels (n=636), (Fig. 1b). This disentanglement between newly synthesized and total RNAs in *ino80*Δ suggests the presence of a compensatory mechanism that buffers mRNA transcription^47^.

The steady state cellular mRNA abundance is the outcome of the opposing actions of mRNA synthesis and degradation. To understand how total mRNA levels remain unchanged, or even increase while transcription rates are reduced in the absence of INO80, we conducted comparative dynamic transcriptome analysis (cDTA)^48^, which measures mRNA synthesis and decay rates (SR and DR respectively). Comparison of the changes in synthesis and decay rates between *ino80*Δ and WT revealed a positive correlation (r=0.47, p-value=2.9e^-233^, Fig 1c). This reduction in mRNA decay rates suggests that mRNA degradation is compromised in *ino80*Δ.

To confirm the stabilization of mRNAs in the absence of INO80, we examined mRNA decay rates of several genes following transcription inhibition with thiolutin. Thiolutin was applied in WT and *ino80*Δ cells at a concentration below the level previously reported to inhibit mRNA degradation^49^ and RNA samples were collected at various timepoints. Following normalization to the stable, Pol III transcribed, *SCR1* RNA, our analysis showed that genes such as *NRD1*, *RNA1*, and *HPT1* exhibited higher mRNA levels in *ino80*Δ compared to WT after transcription inhibition (Fig. 1d and Extended Data Fig. 1b, c). Therefore, our findings indicate that RNA degradation is defective in the absence of INO80 and link the effect of INO80 loss in transcription to changes in RNA decay.

### Loss of INO80 disrupts Pol II progression during transcription elongation

To gain insights into how INO80 promotes transcription, we conducted Native Elongating Transcript coupled to sequencing (NET-seq) analysis in WT and *ino80*Δ yeast cells (Extended Data Fig. 2a). ΝΕΤ-seq allowed us to examine RNA transcripts associated with transcriptionally engaged Pol II by 3’-end sequencing, thus providing a detailed map of elongating Pol II across the genome at nucleotide resolution^50^. To avoid confusion with the term “nascent RNAs” that is used to describe newly synthesized RNAs that are not necessarily associated with Pol II^51^, we will henceforth refer to the Pol II-associated transcripts captured by NET-seq as Pol II density.

We compared the total Pol II density in mRNA genes in WT and *ino80*Δ cells after normalization by DESeq and removal of duplicated reads (Fig. 2a and Extended Data Fig. 2b). 4% of mRNA genes exhibited increased total Pol II density in *ino80*Δ, while only 6% showed significantly decreased Pol II density compared to WT (Fig. 2a). This indicates no substantial changes in overall Pol II occupancy at genes, suggesting that the reduction in transcription output detected by 4tU-seq in *ino80*Δ is not associated with altered transcription initiation.

**Fig. 2.**
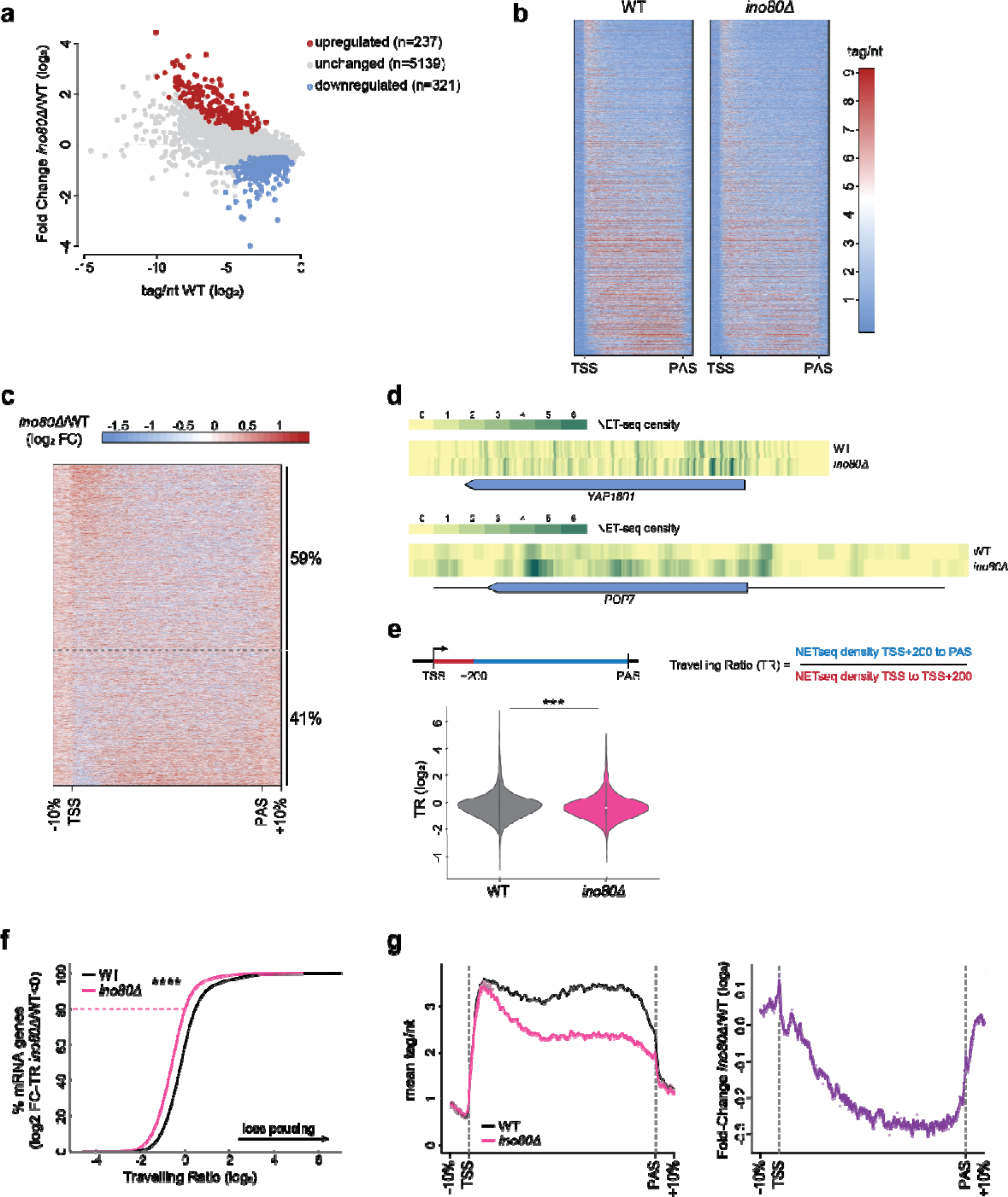
Aberrant Pol II accumulation across the gene body in the absence of INO80 alters Pol II processivity. **a,** Scatterplot analysis for fold changes in NET-seq read density between WT and *ino80*Δ across mRNA genes plotted on a log_2_ scale. Thresholds of 1.5-fold change and 0.05 p values were considered for significant change. **b,** Heatmap analysis of NET-seq read density across mRNA genes (n=4871?) in WT and *ino80*Δ. Tags were aligned to both Transcription Start Site (TSS) and Polyadenylation Site (PAS) for each gene. Genes are sorted by traveling ratio, discussed below. Gradient red-to-blue indicates high-to-low read counts in the corresponding region. **c,** Fold change heatmap analysis comparing the NET-seq read densities in *ino80*Δ and WT across mRNA genes. Genes are listed from lower to higher change in the Traveling Ratio Index between *ino80*Δ and WT (FC-TR *ino80*Δ/WT). Genes above dotted line have decreased Traveling Ratio in *ino80*Δ compared to WT. **d,** Genome browser snapshots of NET-seq read densities across the *YAP1801* and *POP7* genes in WT and *ino80*Δ. Blue coloured box designates the coding sequence. **e,** Left, schematic representation and formula for calculating the Traveling Ratio index (TR) from NET-seq mRNA densities. Right, violin plot of Traveling Ratio values for mRNA genes from a (n=4831). ***p < 0.001 with Mann-Whitney U test. **f,** Cumulative distribution plots comparing the Traveling Ratio (TR) values in WT and *ino80*Δ for mRNA genes with decreased Traveling Ratio in the absence of INO80 (log2 FC-TR *ino80*Δ*/*WT<0). ****p < 0.0001 with Mann-Whitney U test. **g,** Left panel: Metagene analysis for NET-seq read densities in WT and *ino80*Δ across mRNA genes with decreased Traveling Ratios in the absence of INO80. Right panel: Metagene analysis for the fold change in NET-seq read densities between *ino80*Δ and WT across mRNA genes with decreased Traveling Ratios in the absence of INO80. Tags were aligned to both Transcription Start Site (TSS) and Polyadenylation Site (PAS) for each gene. Shaded areas indicate confidence intervals.

The inconsistency between reduced mRNA synthesis and insubstantial decrease in Pol II density in *ino80*Δ prompted us to test whether the distribution of Pol II density across gene units is altered in the absence of INO80 (Fig. 2b). To avoid any potential skew in the analysis from incomplete read coverage we filtered out the bottom 10% of the least transcribed genes. Consistent with previous reports^50,52^ we observed that the density of elongating Pol II was higher at the promoter-proximal region relative to the gene body region for the majority of the genes, indicating that Pol II accumulates downstream of yeast promoters (Fig.2b and Extended Data Fig.2c). Interestingly, deletion of *INO80* appeared to cause a larger decrease in the density of Pol II at the body of the gene compared to the promoter proximal region on average (Extended Data Fig.2c). Fold change analysis for NET-seq read density between *ino80*Δ and WT across gene body regions showed that in the absence of INO80 Pol II density is increased either near the start of the gene at the promoter-proximal region or in downstream areas of the gene body relative to WT (Fig. 2c). Interestingly, the increased accumulation of Pol II at specific gene regions appeared to be associated with a concomitant decrease in Pol II density in other parts of the same gene (Fig. 2c, d and Extended Data Fig. 2d). This suggests that Pol II processivity is altered in the absence of INO80.

To evaluate this possibility, we conducted Traveling Ratio (TR) index analysis, which is calculated as the ratio of Pol II density in the gene body relative to the Pol II density in the first 200bp downstream of the transcription start site (Fig. 2e, scheme). We observed that the Traveling Ratio index for approximately 59% (n=2851) of the genes tested was significantly decreased in *ino80*Δ compared to WT (log2 FC-TR<0), with 525 genes showing log2 fold-change values lower than -1 standard deviation from the average in *ino80*Δ (Fig. 2e, f). In the absence of INO80 Pol II accumulated at the promoter-proximal region relative to the gene body, indicating that Pol II processivity is compromised due to to enhanced promoter proximal pausing (Fig. 2g). Similar results were observed when the bottom 10% of transcribing genes were included in the analysis (data not shown). Contrary to *ino80*Δ, TR index analysis of published NET-seq data^50^ for cells lacking the Rco1 subunit of the gene body specific histone deacetylase Rpd3S complex^53^ showed an increase in Pol II processivity, as expected^54,55^ (Extended Data Fig. 2de). For the remaining 41% of genes in which the Traveling Ratio is not decreased in *ino80*Δ, we observed a higher Pol II density within the gene body compared to the promoter proximal region (Extended Data Fig. 2f, g). These results indicate that loss of INO80 leads to aberrant accumulation of elongating Pol II at specific regions across the gene body and deregulated processivity.

### Loss of INO80 enhances Pol II promoter-proximal stalling

We focused our study on understanding the role of INO80 in counteracting the accumulation of Pol II at the promoter proximal region. Stalling or arrest of Pol II during transcription elongation disrupts Pol II processivity and can alter the traveling ratio of a gene. This prompted us to explore whether INO80 controls Pol II stalling within genes. Several sequences have been reported at which Pol II tends to stall^56^. In *S.cerevisiae* the most enriched logo associated with Pol II pausing within mRNA genes is the ATG motif, where Pol II increasingly stalls after transcribing the A_-1_ and before the T_+1_ positions^50,56^. We aligned the NET-seq data to the A nucleotide of the ATG motif and analysed the Pol II density at the ΑΤG sequences that are found either at the first 200bp from TSS or the downstream gene body region (Fig. 3a and Extended Data Fig. 3a, b). Interestingly, we observed that Pol II density is higher at the promoter proximal ATG sites compared to the ATG sites in the gene body (Fig. 3a), indicating increased stalling early during elongation. Pol II accumulation observed at ATG sites was specific, as no increase was observed when Pol II position was centred at random non-ATG sequences in promoter-proximal regions (Extended Data Fig. 3c). Pol II accumulation at promoter-proximal ATG (AUG) sites was also observed in analysis on published Rpb1 CRAC data^57^, albeit with lower resolution (Extended Data Fig 3d). Analysis of published NET-seq data^50^ at ATG sites showed that deletion of *RCO1* led to decreased accumulation of Pol II at the A nucleotide, while the Pol II pausing profile was altered in cells deleted for the transcription elongation and RNA cleavage stimulating factor Dst1, while (Extended Data Fig. 3e, f). These results corroborate previous findings that Pol II pausing is associated with backtracking and chromatin structure^50^. Strikingly, deletion of INO80 led to a significant increase in Pol II density at promoter-proximal ATG sites, while the change in Pol II density at the gene body ATG sites was less pronounced (Fig. 3b, c). This indicates increased accumulation of stalled Pol II proximally to promoters in the absence of INO80.

**Fig. 3.**
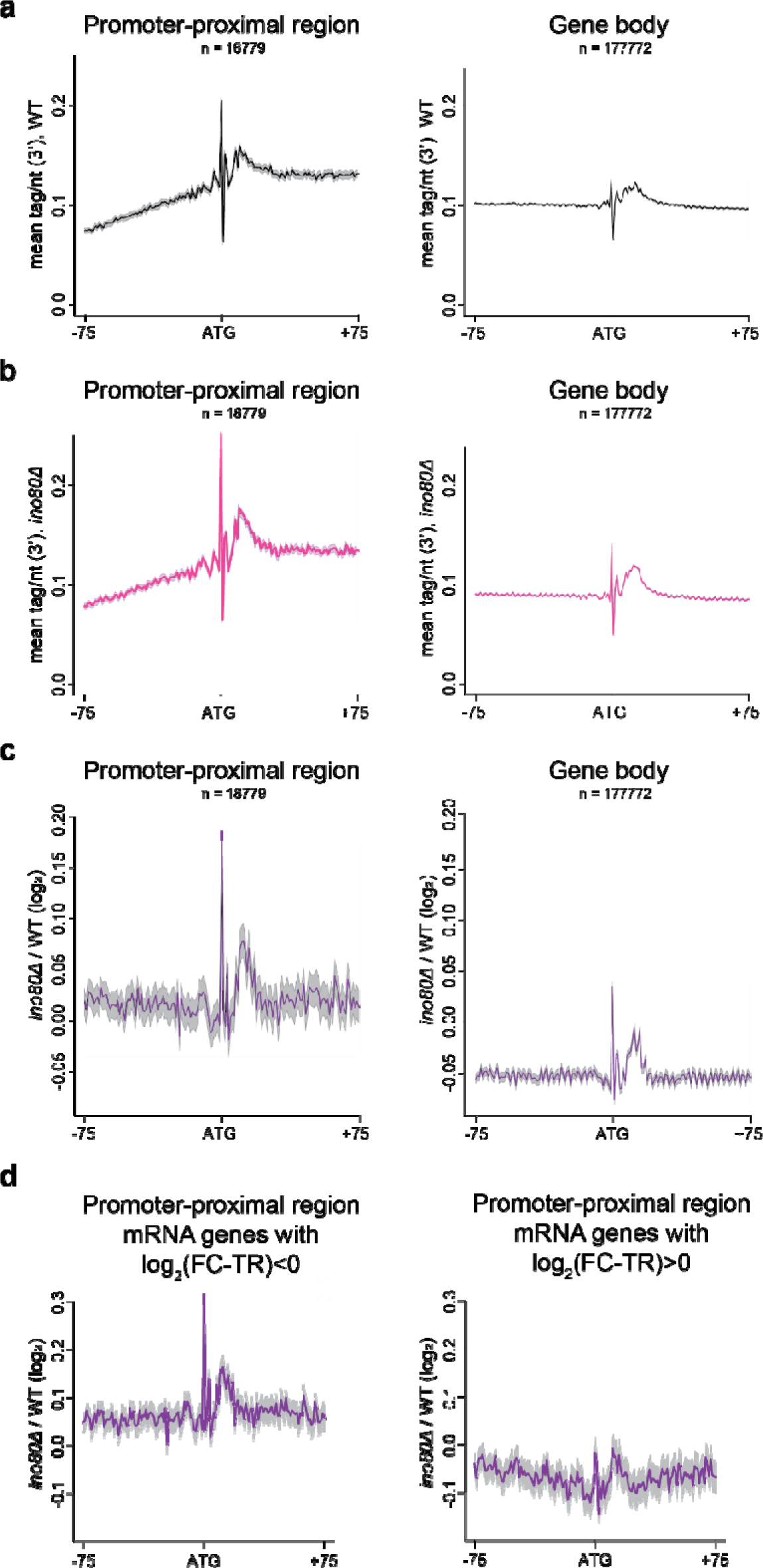
Increased accumulation of Pol II at DNA sequence-determined pausing sites in *ino80*Δ. **a, b,** Metagene analysis for NET-seq read densities in WT (a) and *ino80*Δ (b) aligned at the A nucleotide of the ATG sites in the promoter-proximal (TSS to TSS+200bp) and the gene body (TSS+200 to PAS) regions. n = number of ATG sites in each region. **c,** Metagene analysis for the fold change in NET-seq densities between WT and *ino80*Δ at the promoter-proximal and gene body ATG sites. The mean of the ratio *ino80*Δ/WT for ATG sites in the two regions was calculated and plotted on a log_2_ scale. **d,** Fold change metagene analysis comparing the NET-seq densities between WT and *ino80*Δ at the promoter-proximal ATG sites in genes with increased (left panel) or decreased (right panel) promoter proximal pausing in *ino80*Δ. Left panel: Fold change NET-seq analysis for genes with FC-TR (log2) *ino80*Δ*/*WT <0. Right panel: Fold change NET-seq analysis for genes with FC-TR (log2) *ino80*Δ*/*WT >0.

We hypothesized that the role of INO80 in promoting Pol II processivity maybe linked to alleviation of stalled Pol II. To test this possibility, we examined the Pol II density at promoter-proximal ATG sequences in genes with increased promoter proximal pausing in *ino80*Δ. We grouped genes based on increased and decreased TR in *ino80*Δ compared to WT (log2-FC TR *ino80*Δ*/*WT<0 and log2-FC TR *ino80*Δ*/*WT>0 respectively) and analysed the change in Pol II density at the promoter-proximal ATGs in the two groups. While genes with increased TR in *ino80*Δ showed no change in Pol II density at the promoter-proximal ATG sites, genes with increased promoter proximal pausing showed increased Pol II accumulation at the promoter-proximal ATG sites in the absence of INO80 (Fig. 3d). This suggests that INO80 promotes Pol II processivity by alleviating stalled Pol II.

### Defective transcription elongation in the absence of INO80

Our finding that Pol II processivity is disrupted in the absence of INO80 led us to investigate whether INO80 affects transcription elongation. While the majority of INO80 has been found to be mapped at the +1 nucleosome, INO80 is also enriched further downstream the gene body^41,58^. We therefore evaluated whether the intragenic recruitment of INO80 correlated with Pol II levels. To ensure that the enrichment of Ino80 and Pol II from the promoter region was not included in our analysis, we employed ChIP-exo, which enables near single-nucleotide resolution of factor binding to DNA^59^. We compared the Ino80^41^ and Rpb1 ChIP-exo densities between the transcription start site (TSS) and the polyadenylation site (PAS), as well as within the gene body region from 200bp downstream of the TSS to the PAS. In both regions, we found a strong, positive correlation between Ino80 and Rpb1 enrichment within the gene body, irrespective of whether the first 200bp of the gene body were included or not (Fig. 4a, r=0.63 and Extended Data Fig. 4a, r=0.68, respectively). This indicates that the recruitment of Ino80 within the gene body is associated with Pol II activity, suggesting that INO80 is involved in transcription elongation.

**Fig. 4.**
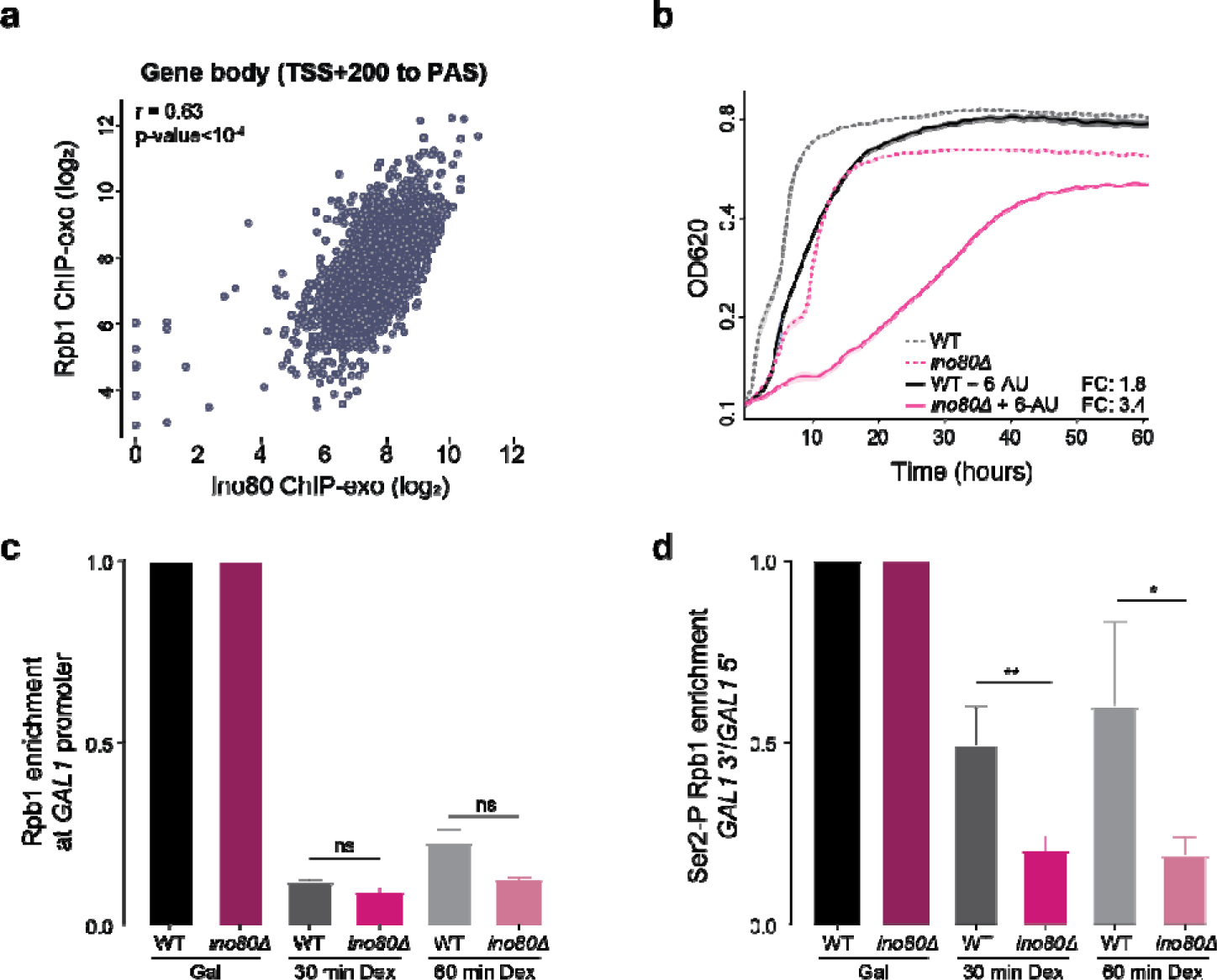
Transcription elongation is defective in the absence of INO80. **a,** Scatterplot analysis comparing Ino80 and Rpb1 ChIP-exo densities between 200bp downstream of transcription start site (TSS+200bp) and the polyadenylation site (PAS) of the bodies of 5798 mRNA genes. Pearson’s correlation coefficient (r) and p-values are indicated. ChIP-exo data for Ino80 derived from ^40^. **b,** Cell proliferation analysis for WT and *ino80*Δ strains grown exponentially in SC-ura and in SC-ura containing 50 µg/ml 6-azauracil (6-AU) liquid media for the indicated time. Cell density was measured at OD^620^ and plotted on a log_2_ scale. Doubling time was calculated from OD^620^: 0.2 to OD^620^: 0.4 for all strains in both conditions. Fold Change ratio (FC) of doubling time in 6-AU over SC-ura for each strain is shown. n=4. **c,** Chromatin immunoprecipitation for Rpb1 at the *GAL1* promoter in WT and *ino80*Δ cells grown in galactose and after 30 and 60 min of dextrose addition in the galactose-containing medium. Fold enrichment at the *GAL1* gene promoter gene was calculated relative to the *INO1* promoter after correction for input DNA levels. The values in dextrose were determined relative to the values in galactose conditions for each strain. Bars, standard errors from three independent experiments. p-values were calculated by t-test after testing for normal distribution by Shapiro-Wilk test. ns; non-significant. **d,** Chromatin immunoprecipitation analysis for the phosphorylated Rpb1 at Serine 2 in WT and *ino80*Δ cells grown in galactose and after 30 and 60 min of dextrose addition in the galactose-containing medium. The *GAL1* 3’/ *GAL1* 5’ ratio was calculated as the Ser2-P Rpb1 enrichment at the 3’ end of the *GAL1* gene relative to the *GAL1* promoter-proximal region (5’). The 3’/5’ ratios in dextrose were determined relative to the values in galactose conditions for each strain. p-values were calculated by t-test after testing for normal distribution by Shapiro-Wilk test. *, p<0.05. **, p<0.01.

We tested whether *ino80*Δ cells are sensitive to transcription elongation stress. We treated cells with 6-azauracil (6-AU), an inhibitor of GTP biosynthesis known to compromise the growth of transcriptional elongation mutants^60^. We observed that *ino80*Δ cells, as well as cells lacking the INO80-specific subunits Arp5 and Arp8, which are crucial for the chromatin remodeling activity and functionality of the complex^38^, grew worse than WT upon 6-AU treatment (Fig. 4b and Extended Data Fig. 4b, c). This sensitivity to 6-AU is consistent with a function for INO80 in transcription elongation.

To test whether INO80 promotes transcription elongation, we analyzed the transcription rates of the galactose-inducible *GAL1* gene. WT and *ino80*Δ cells were grown in galactose and subsequently transferred to dextrose to repress the loading of Polymerase at the *GAL1* promoter. By conducting Rpb1 ChIP, we observed a comparable decrease in the Pol II levels at the *GAL1* promoter in both strains, indicating effective transcriptional shutdown during the experimental time course (Fig. 4c).

To evaluate how Pol II transverses inside the gene body in the absence of INO80 we analyzed the enrichment of the elongating form of Pol II phosphorylated at Serine 2 (Ser2-P Rpb1) at the 3’ and 5’ ends of the *GAL1* gene in galactose and after dextrose addition. Comparison of the 3’/5’ ratios of Ser2-P Rpb1 before and after transcriptional repression determines the extent to which elongating Pol II reaches the gene’s 3’ end, providing an estimate of transcription elongation rates^61^. We observed that when *ino80*Δ cells were shifted to dextrose, the ratio of Ser2-P Rpb1 enrichment at the 3’ end relative to the 5’ end of the *GAL1* gene was significantly reduced compared to WT (Fig. 4d). This indicates that progression of Pol II from the promoter-proximal region to the end of the gene is defective in the absence of INO80 and suggests that relieve of paused Pol II at the promoter proximal region by INO80 promotes transcription elongation.

### Promoter-proximal pausing is associated with premature transcription termination

We sought to understand how INO80 relieves accumulation of stalled Polymerase near the promoters. Previous studies in metazoans have shown that promoter-proximal paused Polymerase is target for premature transcription termination^6,31,32^. We therefore questioned whether promoter-proximal accumulation of Pol II in yeast is associated with RNA surveillance and premature termination. Multiple components of the NNS/Rrp6 non-coding RNA control pathway, such as the nuclear RNA surveillance factor Nab2, the NNS subunit and termination factor Nab3, and the TRAMP subunit Mtr4 have been reported to bind transcripts terminated near the promoters^4^. To explore the link between Pol II pausing and the non-coding RNA control pathway, we calculated the CRAC RNA binding for Nab2, Mtr4, and Nab3^4,57^ in either the promoter-proximal region (TSS to TSS+200bp) or the gene body region (TSS+200 to PAS) of the mRNA transcript, and normalized each value to the transcription levels of the corresponding gene. This allowed us to generate a Transcript Instability index for either the promoter-proximal region (TI^prom^), or the gene body (TI^gene^), which reflects the relative levels of premature termination occurring in the two different regions in each gene (Fig. 5a). To analyse the different TIs in respect to Pol II pausing, we grouped mRNA genes into four clusters based on their TR values in WT conditions. Cluster 1 displaying genes with the lowest TR values hence higher Pol II promoter proximal pausing, while cluster IV included genes with the highest TR values, indicating very little, or no promoter-proximal pausing (Fig. 5b). Strikingly, analysis of the two TIs in the four gene groups revealed a reciprocal relationship between TR and promoter-proximal TI (TI^prom^) for all Nab2, Mtr4, and Nab3 proteins (Fig. 5c): genes with high promoter proximal pausing and low TR exhibited increased promoter-proximal RNA binding of Mtr4, Nab3, and Nab2 (Fig. 5c). Contrary, genes with little or no promoter-proximal accumulation of Pol II (high TR) showed low promoter-proximal TI for the three RNA factors (Fig. 5c). No significant change in the gene body TI values (TI^gene^) for Nab2, Mtr4 and Nab3 was observed among the four TR groups (Fig. 5c). This result indicates that the reduced processivity of Pol II correlates with the association of termination factors in the same RNA region, linking promoter-proximal polymerase pausing with non-coding RNA surveillance and transcription termination.

**Fig. 5.**
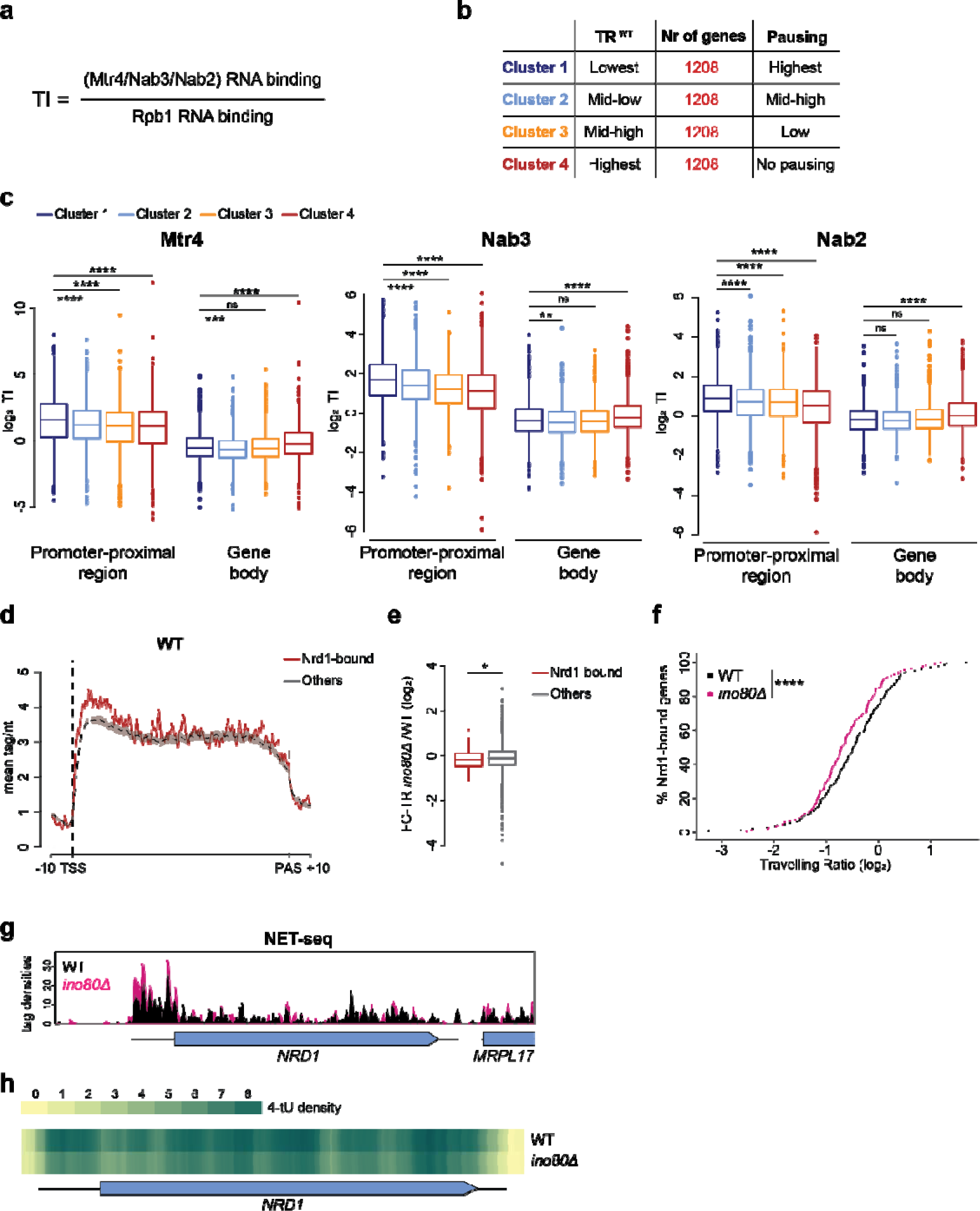
Loading of the RNA surveillance and termination factors Mtr4, Nab3 and Nab2 on promoter-proximal mRNA transcripts is associated with Pol II promoter-proximal pausing. **a,** Formula for calculating Transcript Instability (TI) as the ratio of the averaged CRAC read densities of Nab3^57^, Mtr4 or Nab2^4^ to CRAC densities of Rpb1^57^ for mRNA genes. Two different TI indexes, one for the promoter proximal region ‘TSS to TSS+200’ and one for the gene body region ‘TSS+200bp to TES’ have been calculated. **b,** Table illustrating clustering of mRNA genes according to the relative TR in WT (TR^WT^). The number of genes and the relative level of promoter-proximal pausing are indicated for each cluster. **c,** Boxplot analysis comparing Transcript Instability values for Mtr4, Nab3 and Nab2 calculated for the promoter-proximal and gene body regions of mRNA genes in the four clusters defined by their relative TR^WT^. p-values were calculated by Wilcoxon rank-sum test. ****, p<0.0001. ***, p<0.001. **, p<0.01. ns, non-significant. **d,** Metagene analysis for NET-seq read densities in 143 mRNA genes whose transcripts are bound by Nrd1 at their promoter proximal region (“Nrd1-bound”) and in the remaining mRNA genes of similar transcription levels designated as “Others”. Tags were aligned to both Transcription Start Site (TSS) and Polyadenylation Site (PAS) for each gene. **e,** Boxplot analysis comparing the fold-change in traveling ratio (FC-TR) between *ino80*Δ and WT for the Nrd1-bound and Others groups of genes. p-value was calculated by Wilcoxon rank-sum test. **f,** Cumulative distribution plot comparing the Traveling Ratio (TR) values among Nrd1-bound mRNA genes in WT and *ino80*Δ. ****p < 0.0001 with Mann-Whitney U test. **g,** Genome browser snapshot comparing the NET-seq read densities in WT and *ino80*Δ across the *NRD1* gene. **h,** Genome browser snapshot comparing the 4tU-seq read densities in WT and *ino80*Δ across the *NRD1* gene.

To provide supporting evidence that non-coding RNA transcription termination defines Pol II accumulation at the promoter-proximal region, we a group of 143 mRNA genes that Nrd1 binds strongly to the promoter-proximal region of their transcripts, as identified by the stringent and high-resolution PAR-CLIP assay^27^, and compared them to the remaining mRNA genes with similar transcription levels, that we designated as “Others”. The Nrd1-bound group is enriched in genes attenuated by the NNS-dependent transcription termination pathway, such as *NRD1* and *URA8* (Extended Data Table 1). As anticipated, binding of Mtr4, Nab2 and Nab3 at the promoter proximal regions of the Nrd1-bound RNA transcripts was significantly higher compared to the binding to the “Others” transcripts (Extended Data Fig. 5a). This confirms that NNS binding to mRNAs is associated with premature termination and increased levels of transcript instability. Notably, our NET-seq analysis indicated that Pol II accumulates at higher levels at the promoter-proximal region of the Nrd1-bound genes compared to the “Others” group of genes (Fig. 5d). This reinforces the idea that premature transcription termination is associated with Polymerase pausing.

Intriguingly, we observed that loss of INO80 led to a significant decrease in the TR of the Nrd1-bound genes, indicating that INO80 alleviates promoter-proximal pausing at NNS-regulated genes (Fig. 5e, f). Strikingly, almost all Nrd1-bound genes showed decreased synthesis rates (SR) and decay rates (DR) in *ino80*Δ cells (91% and 98% respectively) suggesting that INO80 controls their expression (Extended Data Fig. 5b and Extended Data Table 1), To explore further the role of INO80 in NNS-regulated genes, we focused on the *NRD1* gene, which is a well-characterized target for promoter-proximal transcription termination by the NNS/Rrp6 pathway^8,62^. We observed by NET-seq that loss of INO80 led to an increase in Pol II density at the *NRD1* promoter-proximal region, but not further downstream (Fig. 5g). Additionally, 4tU-seq analysis showed that the *NRD1* RNA synthesis rate is decreased in *ino80*Δ by more than two-fold compared to WT (Fig. 5h). The inverse relationship between greater promoter proximal pausing and decreased RNA synthesis suggests that Pol II arrests at the *NRD1* promoter proximal termination region in the absence of INO80, leading to downregulation of *NRD1* expression. Taken together, our results indicate that increased promoter proximal stalling in genes regulated by the NNS complex in the absence of INO80 is associated with reduced gene expression.

### Functional and physical interactions between INO80 and non-coding transcription termination factors

Our findings indicate that INO80’s role in relieving stalled Pol II is closely linked to non-coding transcription termination. To investigate the functional connection between INO80 and the NNS-dependent non-coding transcription termination pathway, we examined the genetic interactions of *ARP8* deletion with the termination-defective mutant *nrd1-V368G* (referred to as *nrd1-5*)^8^. Cells lacking *ARP8* were moderately sensitivity to transcription elongation stress induced by 6-AU (Fig. 6a) Contrary, *nrd1-5* mutant cells showed no sensitivity to 6-AU (Fig. 6a). Strikingly, deletion of *ARP8* in the *nrd1-5* mutant strain, led to extreme sensitivity and synthetic lethality in the presence of 6-AU (Fig. 6a). This suggests that INO80 and the NNS complex may function in the same pathway.

**Fig. 6.**
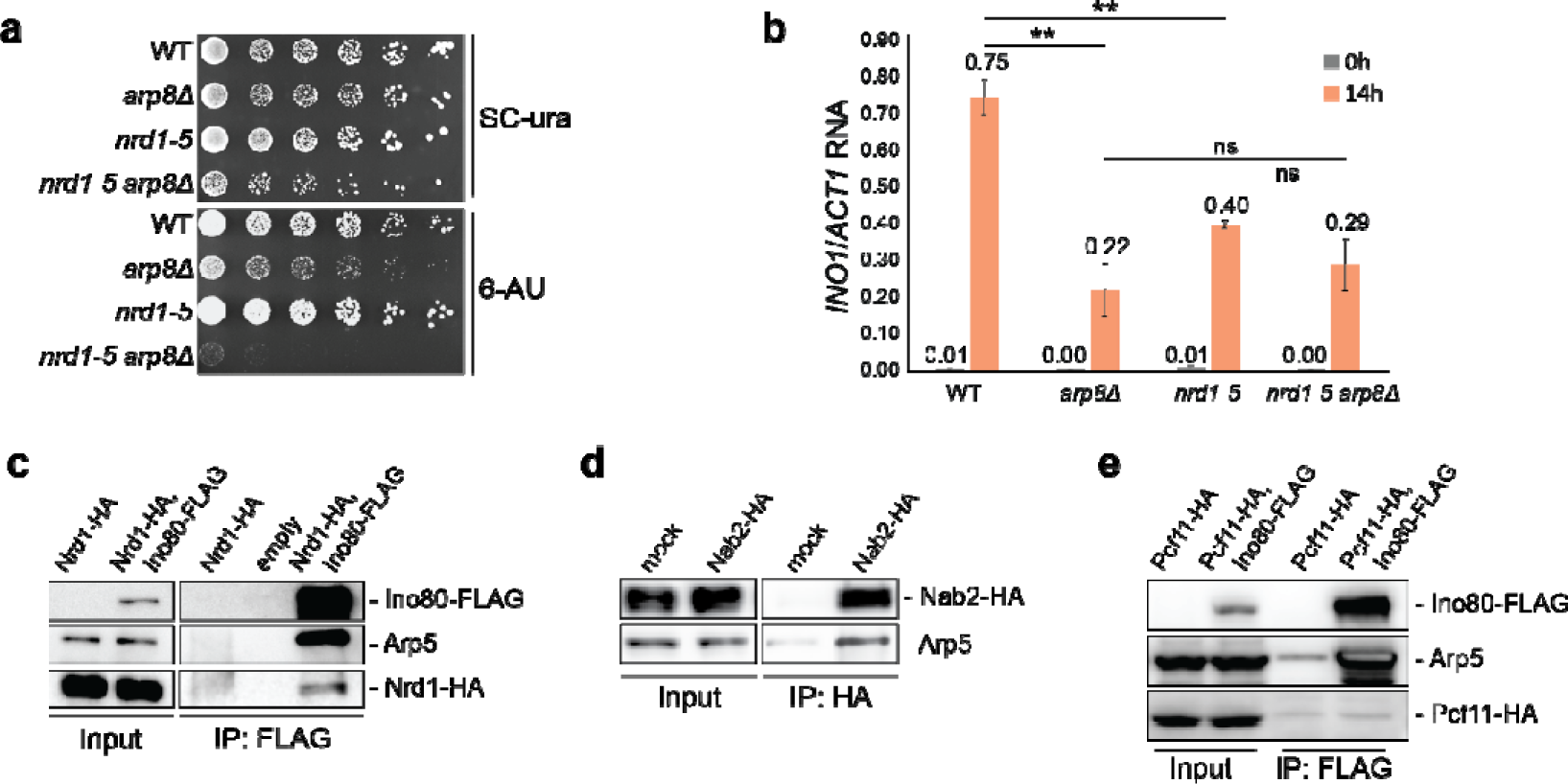
Genetic and physical interactions of INO80 with the non-coding RNA quality control pathway. **a,** 5-fold serial dilution of exponentially grown cells from the indicated strains were plated onto both SC-ura and SC-ura + 50µg/ml 6-Azauracil (6-AU) and incubated at 30°C for 4-5 days. **b,** RT-qPCR analysis for *INO1* RNA was conducted in the indicated strains grown exponentially in SC media (0h) or SC without inositol for 14h. *INO1* RNA was normalized over *ACT1* RNA. Bars, standard errors (n=3). Asterisks indicate statistical significance of the indicated change as calculated by Tukey’s multiple comparisons test. ns: non-significant. **c,** Nucleic acid-free lysates from cells co-expressing Nrd1-HA with Ino80 either untagged or tagged with FLAG epitope were subjected to FLAG-IP. Inputs and IP samples were immunoblotted for FLAG, Arp5, HA. Immunoblot against Pgk1 served as a control. **d,** Nucleic acid-free lysates from cells expressing Nab2-HA were subjected to either HA-IP (Nab2-HA) or IgG-IP (mock). Inputs and IP samples were immunoblotted for Arp5 and HA. **e,** Nucleic acid-free lysates from cells co-expressing Pcf11-HA with Ino80 either untagged or tagged with FLAG epitope were subjected to FLAG-IP. Inputs and IP samples were immunoblotted for FLAG, Arp5 and HA. Samples were resolved in the same SDS-page gel.

The INO80 complex is required for expression of the *INO1* gene in the absence of inositol^63^. We observed that the expression of *INO1* was similarly reduced in both the *arp8* and the *nrd1-5* single mutant strains (Fig. 6b). However, the deletion of *ARP8* in the *nrd1-5* mutant did not further decrease *INO1* expression, indicating an epistatic relationship between the *arp8*Δ and *nrd1-5* mutants (Fig. 6b).

To uncover further evidence linking INO80 to non-coding RNA processing, we analysed published high throughput quantitative genetic interaction data for *INO80* from the CellMap repository^64^ (https://thecellmap.org/). Strikingly, we observed that *INO80* is functionally placed within the non-coding RNA Processing pathway along with Rrp6, Mtr4, and other ncRNA quality control factors (Ext Data Fig. 6a). Similar results from the CellMap repository were obtained for the INO80 specific subunits Ies1, Ies2, Ies3, Ies5 and Nhp10 (data not shown). Contrary, other ATP-dependent chromatin remodeling enzyme genes, such as *SNF2* and SWR1 and General Transcription Factors, such as *TAF1*, *MED11* and *KIN28* are positioned instead in the pathways of Chromatin, Transcription and DNA replication/repair, setting them distinctly apart from the INO80 complex (Ext Data Fig. 6). This functional integration within the non-coding RNA Processing pathway corroborates our genetic results and supports the notion that the INO80 complex operates together with the NNS complex in a coordinated manner.

The functional observations led us to explore whether INO80 physically associates with the transcription termination and RNA quality control machineries. We tested the interactions of INO80 with Nrd1 and the RNA processing factor Nab2, as both are recruited to genes^23,24^. Protein co-immunoprecipitation analysis after nucleic acid removal demonstrated that the INO80 complex physically interacts with both Nrd1 and Nab2 *in vivo* (Fig. 6c, d). However, we did not detect a physical interaction between Ino80 and Pcf11, a component of the cleavage and polyadenylation factor IA complex (Fig. 6e). This lack of association between INO80 and Pcf11 is evident even though INO80 and Pcf11 are both recruited at the 3’ end of mRNA genes and having a physical association with the elongating RNA polymerase II^23,41,43,65^. This *in vivo* analysis indicates that INO80 selectively interacts with machineries involved in the processing of non-coding RNA, implicating INO80 in non-coding RNA transcription termination.

### Non-coding transcription termination is defective in the absence of INO80

To understand the role of INO80 in non-coding RNA transcription termination, we focused on the non-coding snoRNA genes, which are terminated by the NNS complex^8,66^. Using paired-end, spiked-in RNA sequencing we compared the enrichment and length of RNA transcript reads that encompass the snoRNA mature transcript end site (mTES) in WT, *ino80*Δ and in cells deleted for the 3’-5’ exonuclease *RRP6*, which is responsible for the trimming and 3’ end maturation of pre-snoRNAs^67^. as control. Tandem snoRNAs and snoRNAs whose mTES is located less than 250bp upstream of mRNA genes were excluded from the analysis to avoid confounding results. We observed that absence of INO80 led to an increase in snoRNA transcripts that were extended beyond the mTES (Fig. 7a, b). Additionally, the snoRNAs expressed in *ino80*Δ are extended further than the snoRNAs observed in the absence of RRP6 (Fig. 7a, b). This result indicates that the increase in extended snoRNAs in *ino80*Δ is not the consequence of stabilization of snoRNA precursors and suggests that snoRNA termination is defective.

**Fig. 7.**
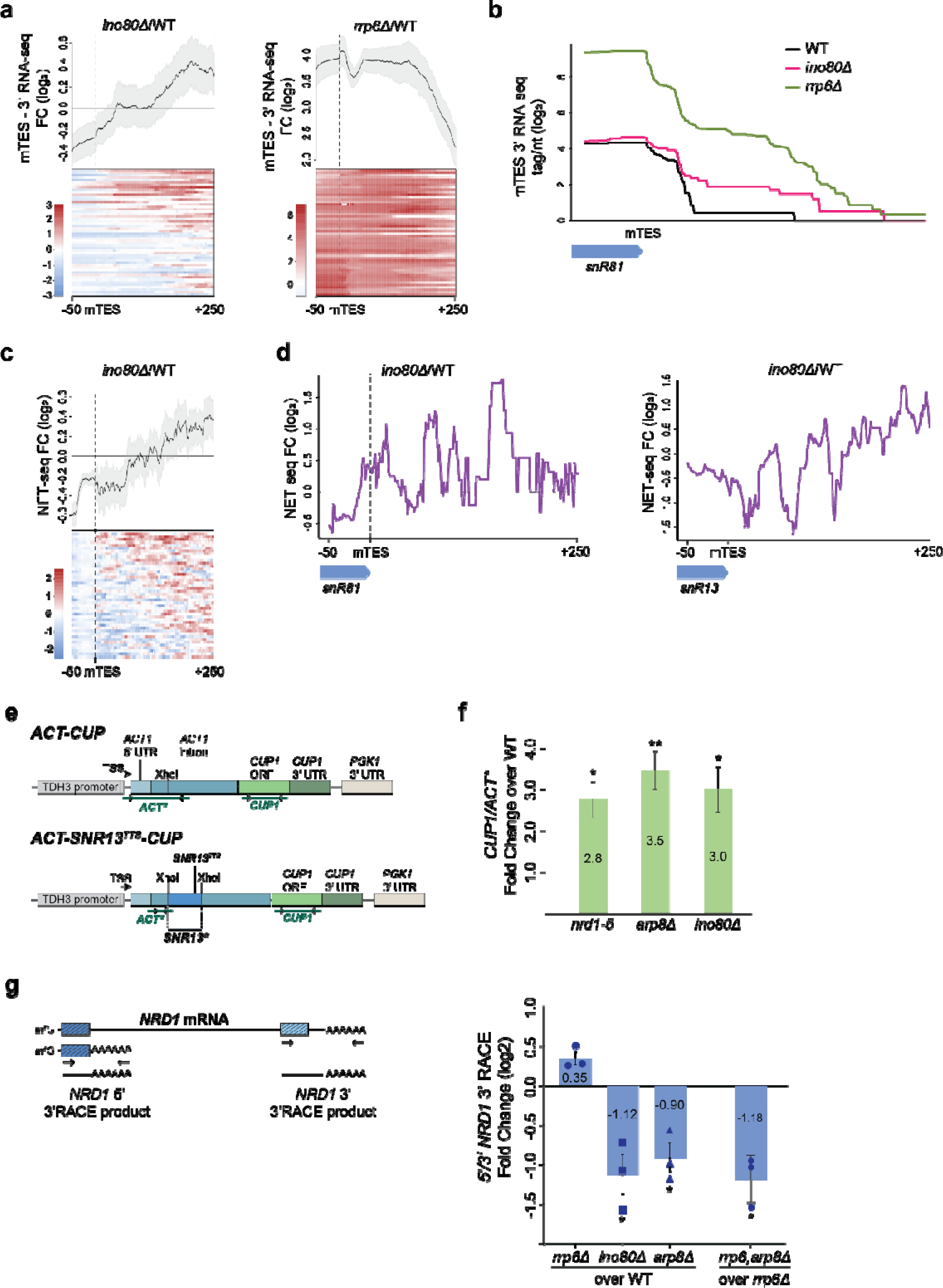
Transcription termination readthrough at non-coding RNA termination sites in the absence of INO80. **a,** Fold change metagene and heatmap analysis comparing RNA-seq read densities at the 3’ end region of snoRNAs between WT and *ino80*Δ (FC *ino80*Δ/WT), and between WT and *rrp6*Δ (FC *rrp6*Δ/WT). Only RNA-seq reads encompassing the mature Transcript End Site (mTES) in each strain were analysed. Tags were aligned to the mTES. Gradient red-to-blue indicates high-to-low read counts in the corresponding region. **b,** Density distribution plot of RNA-seq reads encompassing the mTES *snR81* region in WT, *ino80*Δ and *rrp6*Δ. **c,** Fold change metagene and heatmap analysis comparing NET-seq densities between WT and *ino80*Δ (FC *ino80*Δ/WT) downstream of snoRNA genes (mTES-50bp to mTES+250bp). Tags were aligned to the mTES. Tandem snoRNAs and snoRNAs whose mTES is located less than 250bp upstream of mRNA genes were excluded from the analysis. Gradient red-to-blue indicates high-to-low read counts in the corresponding region. **d,** Fold change density distribution plot comparing NET-seq reads downstream the *SNR81* and *SNR13* genes in WT and *ino80*Δ (FC *ino80*Δ/WT). **e,** Schematic illustration of the *ACT-CUP* and *ACT-SNR^TTS^-CUP* reporter plasmids. The ACT and CUP transcript regions analysed by RT-qPCR are indicated. **f,** RT-qPCR analysis was conducted in WT, *nrd1-5*, *arp8*Δ and *ino80*Δ cells transformed with either *ACT-CUP* or *ACT-SNR^TTS^-CUP* plasmids. The *CUP1*/*ACT\** ratio from the *ACT-SNR^TTS^-CUP* plasmid was normalized over the same ratio from the control *ACT-CUP* plasmid (Extended Data Fig. 5e) to correct for differential transcription across all strains. Read-through at the *SNR13^TTS^* for each indicated strain is shown as Fold Change Ratio over WT. Bars, standard errors from three independent experiments. p-values were calculated by t-test after testing for normal distribution by Shapiro-Wilk test. *, p<0.05. **, p<0.01. **g,** Left: Scheme illustrating nested 3’RACE for termination events at the promoter-proximal (5’) and PAS (3’) regions of the *NRD1* gene (*NRD1 5’*, dark blue). Right: 3’RACE analysis for termination events at the promoter-proximal (5’) and PAS (3’) regions of the *NRD1* gene in WT and the indicated mutant strains. The relative enrichment of polyadenylated promoter-proximal to PAS-terminated transcripts (5’/3’) was calculated in each strain. The fold change 5’/3’ ratios for the single *rrp6*Δ, *ino80*Δ and *arp8*Δ mutants relative to WT and for the *arp8*Δ*,rrp6*Δ double mutant relative to *rrp6*Δ single mutant were analysed and plotted on a log_2_ scale. n=3. p-values were calculated by t-test after testing for normal distribution by Shapiro-Wilk test. *, p<0.05.

INO80 deletion did not affect termination of protein-coding genes such as *RPS7B* (Extended Data Fig. 7a). Furthermore, the expression of known transcription termination and RNA surveillance factors was not significantly altered by the deletion of *INO80* or *ARP8* (Extended Data Table 2). This excludes the possibility that the increase in extended snoRNA transcripts in *ino80*Δ is due to compromised expression of RNA processing and quality control factors.

To test whether INO80 promotes non-coding RNA transcription termination, as suggested by our RNA-seq data analysis, we analyzed Pol II density downstream the snoRNA genes by NET-seq. We observed that, although Pol II inside most snoRNA genes, such as *SNR81* and *SNR13,* was reduced in *ino80*Δ, its density downstream was increased compared to WT (Fig. 7c, d). This indicates that non-coding transcription termination is defective in the absence of INO80.

To evaluate this possibility, we utilized the *ACT-SNR^TTS^-CUP* reporter system in which the insertion of the Transcription Termination Site (TTS) of the snoRNA gene *SNR13* (*SNR^TTS^*) upstream of the *CUP1* gene silences its expression (Fig. 7e)^8,20^. Ino80 and Nrd1 are co-enriched at the endogenous *SNR13^TTS^*region confirming the suitability of the *ACT-SNR^TTS^-CUP* reporter system for investigating the role of INO80 in NNS-dependent transcription termination (Extended Data Fig. 7b). WT cells transformed with the *ACT-SNR^TTS^-CUP* plasmid were unable to grow in copper-containing media due to silencing of the *CUP1* gene (Extended Data Fig. 7c). Contrary, *ino80*Δ cells transformed with *ACT-SNR^TTS^-CUP* grew in the presence of copper, suggesting read-through transcription of *CUP1* (Extended Data Fig. 7c). To monitor *CUP1* transcription, we conducted RT-qPCR analysis using primer pairs targeting regions before and after the *SNR13^TTS^* insertion site in the *ACT-CUP* reporter plasmids with or without the *SNR13^TTS^*element (Fig. 7e). The *nrd1-5* mutant exhibited a 3-fold increase in relative *CUP1* expression from the *ACT-SNR^TTS^-CUP* reporter compared to WT, consistent with previous studies^8^ (Fig. 7f and Extended Data Fig. 7d). The *ino80*Δ and *arp8*Δ mutants also showed a significant increase in relative *CUP1* transcription, with fold changes of 2.8 and 3.5, respectively (Fig. 7f and Extended Data Fig. 7d). Moreover, inducible, acute loss of the Ino80 protein using the *INO80-td* degron system^68^ resulted in a significant increase in relative *CUP1* levels from the *ACT-SNR^TTS^-CUP* plasmid, but not from the control *ACT-CUP* plasmid lacking the transcription termination site (Extended Data Fig. 7e). These results collectively indicate that INO80 promotes non-coding transcription termination.

The effect of INO80 deletion in snoRNA transcription termination prompted us to explore whether INO80 is also involved in NNS-dependent premature termination of mRNA synthesis. Metagene analysis of ChIP-exo data for Ino80^41^ in the Nrd1-bound and “Others” groups of genes demonstrated that Ino80 enrichment extends beyond the transcription start site (TSS) into the promoter-proximal region of Nrd1-bound genes (Extended Data Fig. 7f). This indicates that INO80 is recruited at premature termination regions in mRNA genes. To evaluate the role of INO80 in premature transcription termination, we employed quantitative 3’ rapid amplification of cDNA ends (3’ RACE) for determining the abundance of terminated, polyadenylated transcripts at the promoter-proximal region and the PAS of the *NRD1* gene (Fig. 7g, scheme). The relative enrichment of prematurely terminated transcripts was calculated as the ratio of promoter-proximal poly(A)-RNAs to PAS-terminated transcripts. Deletion of *RRP6* led to a substantial increase in the abundance of both promoter-proximal and full length *NRD1* polyadenylated species, consistent with RNA-seq results and previous studies^62^ (Fig. 7g and Extended Data Fig. 7g, h). On the contrary, deletion of *INO80* or *ARP8* resulted in a significant reduction in the enrichment of promoter-proximal poly(A) transcripts (Fig. 7g). The decrease in promoter-proximal poly(A) transcripts observed in *arp8*Δ was not rescued by deletion of *RRP6* (Fig. 7g), indicating that the reduction in promoter proximal poly(A) transcripts is not due to enhanced RNA degradation. These results indicate defective premature transcription termination of *NRD1* in the absence of INO80 and suggest that the INO80 complex promotes premature transcription termination of mRNA genes.

### Chromatin recruitment of Nab2 and release of Pol II from a non-coding transcription termination site depend on INO80

During non-coding transcription termination, the recruitment of RNA processing factors to chromatin and binding to nascent RNA facilitate the degradation of aberrant RNA by the exosome and dissociation of the terminated Pol II complex from chromatin^21^. We hypothesized that INO80 promotes non-coding transcription termination by enabling the recruitment of the ncRNA processing machinery to chromatin. Nab2 is co-transcriptionally recruited to the gene body of active genes^24^ and binds to prematurely terminated mRNAs^4^, including promoter-proximal *NRD1* and *IMD2* transcripts (Extended Data Fig. 8a). ChIP analysis showed that loss of INO80 resulted in reduced recruitment of Nab2 at both the *NRD1* and *IMD2* promoter proximal regions, despite no decrease in Pol II levels in the same regions (Fig. 8a, Extended Data Fig. 8b-d and Fig. 5g). Likewise, Nab2 recruitment at the *SNR13* transcription termination region was significantly reduced in *ino80*Δ cells compared to WT (Fig. 8b). This is in line with our observation that Pol II density is increased downstream the *SNR13* gene in the absence of INO80 (Fig. 7d). These results indicate that INO80 promotes the recruitment of Nab2 to non-coding transcription termination sites. Co-immunoprecipitation analysis for Nab2 demonstrated that the physical interaction between Nab2 and Rpb1 was unaffected by the loss of INO80 (Extended Data Fig. 8e), ruling out the possibility that the defect in Nab2 recruitment to chromatin is due to compromised binding to the Pol II machinery.

**Fig. 8.**
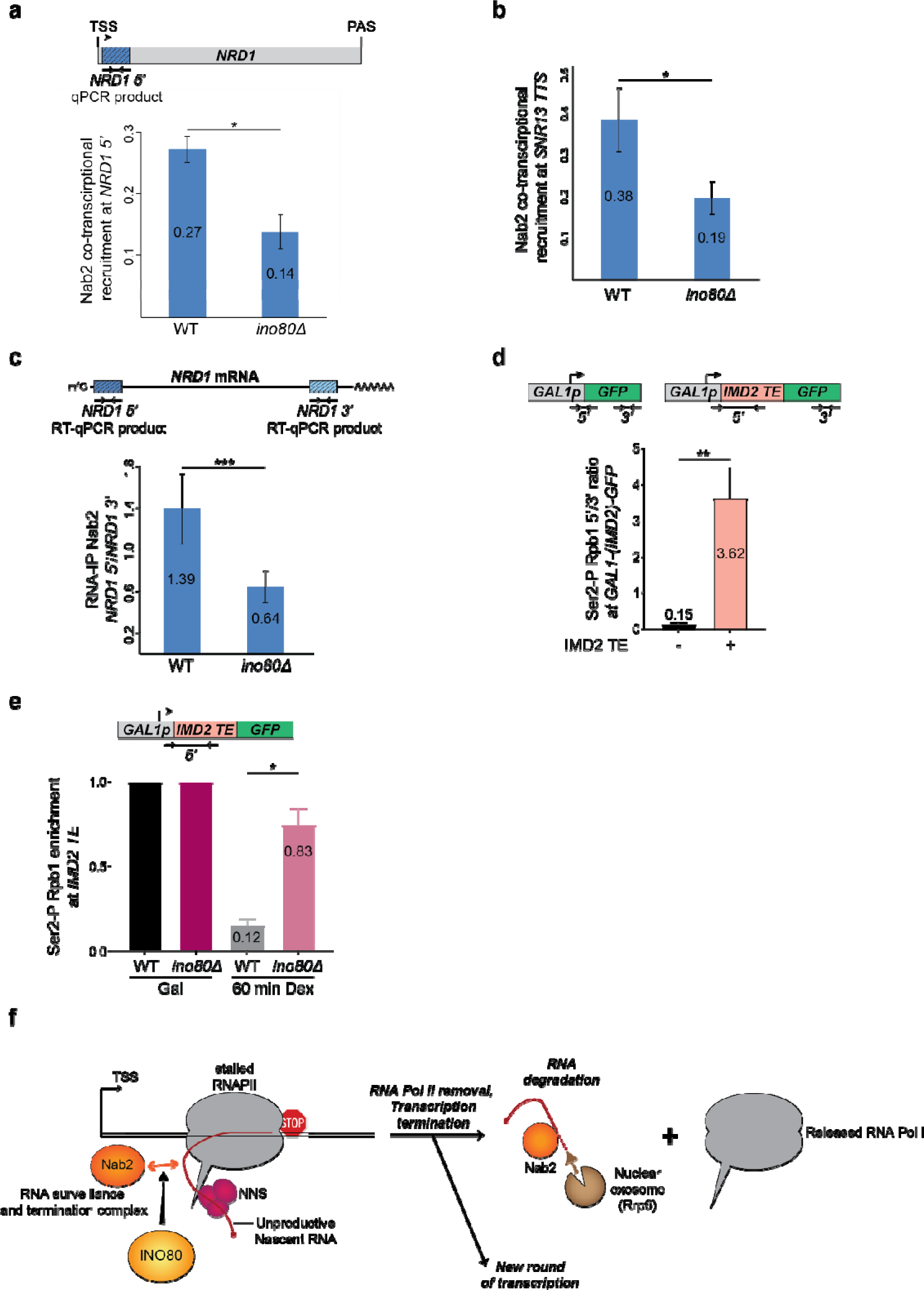
INO80 promotes the co-transcriptional recruitment of Nab2 and the release of stalled Pol II from non-coding transcription termination sites. **a,** Co-transcriptional Nab2 recruitment at the promoter-proximal region of the *NRD1* gene (*NRD1 5’*). Chromatin immunoprecipitation for Nab2-HA and Rpb1 was conducted in WT and *ino80*Δ cells. Fold enrichment for Nab2 and Rpb1 at the *NRD1 5’* (dark blue region in scheme) was calculated relative to the *GAL1* gene after correction for input DNA levels. Co-transcriptional Nab2 recruitment in the two strains was calculated as the ratio of Nab2 over Rpb1 ChIP enrichment in the *NRD1 5’* region. Bars, standard errors from four independent experiments. p-values were calculated by t-test after testing for normal distribution by Shapiro-Wilk test. *, p<0.05. **b,** Co-transcriptional Nab2 recruitment at the Transcription Termination region (TTR) of the *SNR13* gene in WT and *ino80*Δ cells. Analysis was conducted as in (a). **c,** RNA immunoprecipitation for Nab2-HA in WT and *ino80*Δ cells. Ratio of binding of Nab2 to the promoter-proximal *NRD1* RNA (upper panel, *NRD1 5’*, dark blue) over binding to the 3’ region of *NRD1* RNA (upper panel, *NRD1 3’,* light blue) after correction for input (total) RNA levels in the indicated strains. Bars, standard errors from four independent experiments. p-values were calculated by t-test after testing for normal distribution by Shapiro-Wilk test. ***, p<0.001. **d,** Upper panel, schematic representation of the *GAL1p-GFP* and *GAL1p-IMD2 TE-GFP* reporter genes. TE: termination element. Arrows denote the 5’ and 3’ regions amplified by qPCR to calculate the 5’/3’ ChIP enrichment ratio for each reporter gene. Lower panel. Chromatin immunoprecipitation analysis for the phosphorylated Rpb1 at Serine 2 in WT cells expressing either the *GAL1p-GFP* or the *GAL1p-IMD2 TE-GFP* reporter gene and grown in galactose conditions. The 5’/3 ratio was calculated as the Ser2-P Rpb1 enrichment at the 5’ end relative to the 3’ end of the *GAL1p*-GFP reporter gene without (-) or with (+) the *IMD2* termination element. **e,** Chromatin immunoprecipitation analysis for the phosphorylated Rpb1 at Serine 2 in WT and *ino80*Δ cells expressing the *GAL1p-IMD2 TE-GFP* reporter gene and grown in galactose and after 60 min of dextrose addition in the galactose-containing medium. Fold enrichment at the *IMD2 TE* region of the reporter gene was calculated relative to the *INO1* gene after correction for input DNA levels. **f,** Model for INO80-dependent premature transcription termination mechanism. See Discussion for details.

The dependency of Nab2 recruitment to chromatin on INO80 led us to investigate whether INO80 promotes binding of Nab2 to RNA. Quantitative RNA immunoprecipitation (RNA-IP) revealed that the binding of Nab2 to the promoter-proximal region of the *NRD1* transcript was significantly reduced in *ino80*Δ cells relative to its binding to the 3’ end (Fig. 8c). This suggests that INO80-mediated recruitment of Nab2 to chromatin enables its binding to nascent RNAs.

We seek to understand the mechanism by which INO80 promotes non-coding transcription termination. INO80 enables Pol II removal from chromatin^43^. Since our NET-seq analysis indicated that Pol II accumulates at non-coding termination sites in the absence of INO80, we investigated whether INO80 facilitates the release of Pol II from DNA during NNS-dependent transcription termination. We utilized a *GFP* reporter system^69^ in which the *IMD2* non-coding termination element (*IMD2^TE^*) was inserted between the galactose-inducible *GAL1* promoter (*GAL1^prom^*) and the *GFP* gene (Fig. 8d, scheme). The *IMD2^TE^* contains multiple NNS-binding sites that act as transcription termination signals, leading to transcriptional attenuation and silencing of *GFP* expression^70^.

Comparative ChIP analysis of Ser2-P Rpb1 in the presence and absence of the *IMD2 ^TE^* under galactose-inducible conditions revealed a 25-fold increase in the accumulation of elongating Pol II at the promoter-proximal region relative to *GFP* when the *IMD2 ^TE^* was inserted (Fig. 8d). This indicates that the *IMD2* non-coding termination element induces the arrest of Pol II, impeding its progression further downstream.

Upon dextrose addition, total Rpb1 was decreased by roughly 5-fold in both WT and *ino80*Δ cells at the 5’ end of the *GAL1^prom^-IMD2^TE^-GFP* reporter, indicating that transcription is efficiently repressed during the experimental time course (Extended Data Fig. 8f). In the WT strain, the accumulation of Ser2-P Rpb1 at *IMD2 ^TE^* dropped by approximately 90% after addition of dextrose (Fig. 8e), indicating that the majority of arrested Pol II is removed from chromatin. However, in *ino80*Δ cells, Ser2-P Rpb1 enrichment at the IMD2 transcription termination region following dextrose repression was 7-fold higher compared to WT, remaining nearly as high as during galactose induction conditions, (Fig. 8e). This suggests that INO80 promotes the release of Pol II from chromatin during non-coding RNA transcription termination.

## DISCUSSION

In this study, we have expanded our understanding of the role of INO80 in gene expression. Through a comprehensive analysis of nascent transcription rates, Pol II dynamics, functional and physical interactions, and reporter assays, we have uncovered a role for INO80 in termination of transcription mediated by the non-coding RNA quality control pathway. Our findings indicate that INO80 plays a crucial role in the release of stalled Pol II from chromatin, thereby enabling non-coding transcription termination. Our data suggest that removal of stalled Pol II by INO80 during premature transcription termination could potentially allow subsequent rounds of transcription to progress along the gene body, enhancing productive mRNA synthesis and improving transcription rates (Figure 8f).

Previous studies using steady-state RNA transcriptomics approaches have suggested that INO80 can play both a positive and a negative role in transcription in a gene-specific or chromatin context-dependent manner^39,65,71^. Here, we employed nascent transcriptome analysis to examine the immediate transcriptional output in the absence of INO80. Our data revealed a global downregulation of Pol II transcription in the absence of INO80, indicating that the role of INO80 is to facilitate, rather than repress transcription. This observation highlights the importance of studying the dynamic process of transcription to fully understand the role of INO80, as well as transcription and epigenetic factors in general. Interestingly, the decrease in mRNA synthesis rates in the absence of INO80 is coupled with stabilisation of mRNA transcripts. This buffering of steady-state mRNA levels has been reported as a counter-mechanism that maintains cellular mRNA abundance in conditions of global transcriptional stress^48,72^. It is thus possible that the accumulation of stabilized transcripts in the absence of INO80 may have led to previous misconceptions of a negative role for INO80 in transcription.

One key aspect of INO80’s function in transcription that was revealed by our study is its impact on Pol II processivity and pausing. For most of the genes, we observed enhanced promoter proximal pausing and reduced progression of Pol II into the gene body in the absence of INO80. This increase in stalled Pol II was confirmed by our single-nucleotide NET-seq analysis at intragenic pausing sites that are defined by the A^-1^T^+1^G motif^50^. This analysis revealed that the dwell time of Pol II at the promoter-proximal ATG pausing sites is prolonged in genes with reduced Pol II processivity in the absence of INO80. As pausing at ATG sites has been associated with Polymerase backtracking and stalling^50^, our results strongly suggest that INO80 alleviates stalled or terminally arrested Polymerases, which if not timely removed can affect Pol II processivity and gene expression. According to a recent study using *ex-vivo* PRO-seq analysis, it was proposed that the depletion of INO80 causes Pol II to pause at alternative sites in a small number of genes^73^. Whether these alternative sites are associated to the ATG pausing is unclear. However, it should be noted that, while NET-seq can capture both actively elongating and stalled Polymerase molecules, PRO-seq is limited in its ability to detect stalled Polymerases^74,75^. This key difference between the two technical approaches could explain the discrepancies in the results between our study and ^73^.

Interestingly, our NET-seq analysis indicated that in several genes, deletion of INO80 can also lead to enhanced accumulation of Pol II not at the promoter proximal region but further inside the gene body. The reason for this late increase in Pol II density downstream the promoter proximal region is unclear. It is unlikely that this change reflects accelerated early elongation rates at the promoter region in the absence of INO80, as this would eventually lead to a buildup of Pol II throughout the gene. An alternative possibility, given that INO80 has been reported to be recruited inside gene bodies^58^ and is also enriched at transcription termination sites^41,65^, is that INO80 counteracts stalling of Pol II at the late steps of transcription elongation. Further analysis is needed to illuminate how and where Pol II accumulates during late transcription in the absence of INO80.

How does Pol II accumulate at pausing sites in the absence of INO80? While it is possible that absence of INO80 makes elongating Pol II more prone to arrest, this explanation is unlikely. Increased frequency of stalling would result in a greater number of polymerases being terminated, as observed in the absence of the cleavage stimulating factor TFIIS/Dst1 for example^76^. This is contrary to what has been observed in the *ino80*Δ mutant in which degradation of Pol II^42,43^ and RNA decay are compromised. Importantly, we observed that release of stalled Polymerase from chromatin is defective in the absence of INO80. This finding further supports a role for INO80 in removing Polymerases after they have been stalled or arrested. Additionally, this finding is consistent with biochemical evidence demonstrating the physical association between INO80 and the Pol II elongation machinery^42,43^ and the recently reported role for INO80 in remodelling of hexasomes, the subnucleosomal particles resulting from Pol II transcription^77^.

Notably, our study uncovered a strong association between Pol II processivity and pausing and premature transcription termination. We observed a clear correlation between the accumulation of paused Pol II at promoter-proximal regions and the binding of non-coding RNA surveillance and termination factors, such as Nab3, Mtr4, and Nab2, to the same RNA domains. This association was particularly prominent in genes that are attenuated by the NNS complex, suggesting a mechanistic link between Pol II pausing and non-coding transcription termination. These observations validate and expand earlier *in vitro* and *in vivo* studies that suggested that slowdown of elongating Pol II can establish a temporal and spatial window for Pol II termination by the non-coding transcription termination pathway^78,79^. While it is possible that promoter-proximal pausing reflects faulty Polymerase complexes unable to move into productive elongation, recent evidence has shown that elongation factors are properly recruited to promoter-proximally paused Pol II, making this scenario less likely^80^. Therefore, our results support a model in which pausing or stalling of Pol II at specific promoter-proximal sites allows for the efficient loading of RNA surveillance and termination factors, leading to the timely termination of transcription. Fascinatingly, several lines of evidence suggest that INO80 connects stalling of Pol II to the non-coding transcription termination machinery: (i) deletion of INO80 resulted in a significant increase in Pol II pausing at NNS attenuated genes with a concomitant decrease in Pol II processivity and transcription rates; (ii) the increased Pol II density observed at the promoter-proximal non-coding termination regions of the *NRD1* and *IMD2* genes in the absence of INO80 was accompanied by compromised recruitment of Nab2 at the same regions and a reduction in prematurely terminated transcripts; (iii) removal of stalled Pol II from the non-coding transcription termination region of the *IMD2* gene is dependent on INO80. Given that loss of INO80 results in globally reduced transcription rates, it is therefore tempting to speculate that by coupling RNA surveillance to transcriptional pausing, chromatin acts as a molecular switch that dictates whether Pol II will either progress into productive elongation or be removed to allow for the next round of transcription to take place, hence improving the overall processivity of Pol II. Consistently, our NET-seq and RNA-seq analyses across snoRNA genes and analysis of the *ACT-SNR^TTS^-CUP1* reporter system demonstrated that INO80 is essential for efficient snoRNA transcription termination. These collective findings highlight a previously uncharacterized role for chromatin remodeling complexes, such as INO80, in orchestrating the precise termination of non-coding RNAs.

How does INO80 regulate non-coding transcription termination? Co-immunoprecipitation analysis identified a physical interaction between the INO80 complex and the transcription termination and RNA surveillance factors Nrd1 and Nab2. Deletion of the INO80-specific subunit *ARP*8 in the *nrd1-V368G* mutant strain led to synthetic lethality upon transcriptional stress conditions, highlighting the cooperative nature of these complexes. Together with the observation that INO80 is enriched at NNS-dependent termination regions, such as the *SNR13* TTS, our results suggest a direct involvement for INO80 in the co-transcriptional non-coding RNA quality control pathway.

A critical aspect of non-coding transcription termination is the release of Pol II from chromatin, which relies on the co-transcriptional binding of termination and surveillance factors to both the nascent RNA and the Pol II machinery^21^. Our study found that INO80 promotes the dissociation of stalled Pol II from non-coding RNA termination sites upon transcriptional shutdown and facilitates the recruitment of Nab2 to chromatin, hence revealing a crucial role for INO80 in the process. This suggests that chromatin remodeling is necessary for the correct formation of ribonucleoprotein particles (RNPs) necessary for the destabilization of the transcription elongation complex and subsequent release of Pol II from chromatin. Therefore, in agreement with our genome-wide data, our molecular analysis using established gene models with well characterized sites of non-coding transcription termination raises the possibility that INO80 enables non-coding transcription termination by promoting the recruitment of RNA surveillance factors to chromatin.

Interestingly, INO80 has been shown to interact with and facilitate the release of poly-ubiquitinated Rpb1 from chromatin^43^ suggesting that Rpb1 ubiquitination could be involved in the process. This raises the intriguing possibility that INO80 selectively evicts Pol II molecules by recognizing and acting on ubiquitinated Rpb1, providing specificity to the termination process.

Recent studies have highlighted the connection between promoter-proximal transcriptional pausing and premature termination in metazoans^6,30–32^, while dysregulated Pol II pausing and uncontrolled transcription elongation have been implicated in human diseases^81,82^. Mutations in the RNA surveillance machinery have also been linked to increased transcriptional stress and genomic instability^83^, both hallmarks of cancer^84^. Notably, dysregulated expression of INO80 has been associated with disease^38,85,86^. Thus, we speculate that the findings presented in this study could offer insights into the molecular mechanisms underlying the involvement of INO80 in disease pathogenesis.

## Acknowledgements

We thank D. Shapira (Newcastle University) for assisting with experiments and constructive discussions. We thank S. Baulande and P. Legoix-Né (NGS platform, Institute Curie). We also thank Maxime Wery and Nicolas Vogt for outstanding help with the NET-seq and RNA-seq assays and critical reading of the manuscript, David A. Brow for the generous gift of the pGAC24 and pGAC24-SNR13TTS plasmids and the *nrd1-5* yeast strain, and Kuangyu Yen, Lisa Prendergast and members of the Papamichos-Chronakis and Morillon’s labs for constructive discussions. This work has benefited from the facilities and expertise of the NGS platform of Institute Curie, supported by the Agence Nationale de la Recherche (ANR-10-EQPX-03, ANR10-INBS-09-08) and the Canceropôle Ile-de-France.

## Data availability statement

Accession numbers: Raw sequences have been deposited to the NCBI Gene Expression Omnibus as Series GSE108046, GSE108053, GSE108054, GSE185733.

## EXTENDED DATA FIGURES

**Extended Data Fig. 1.**
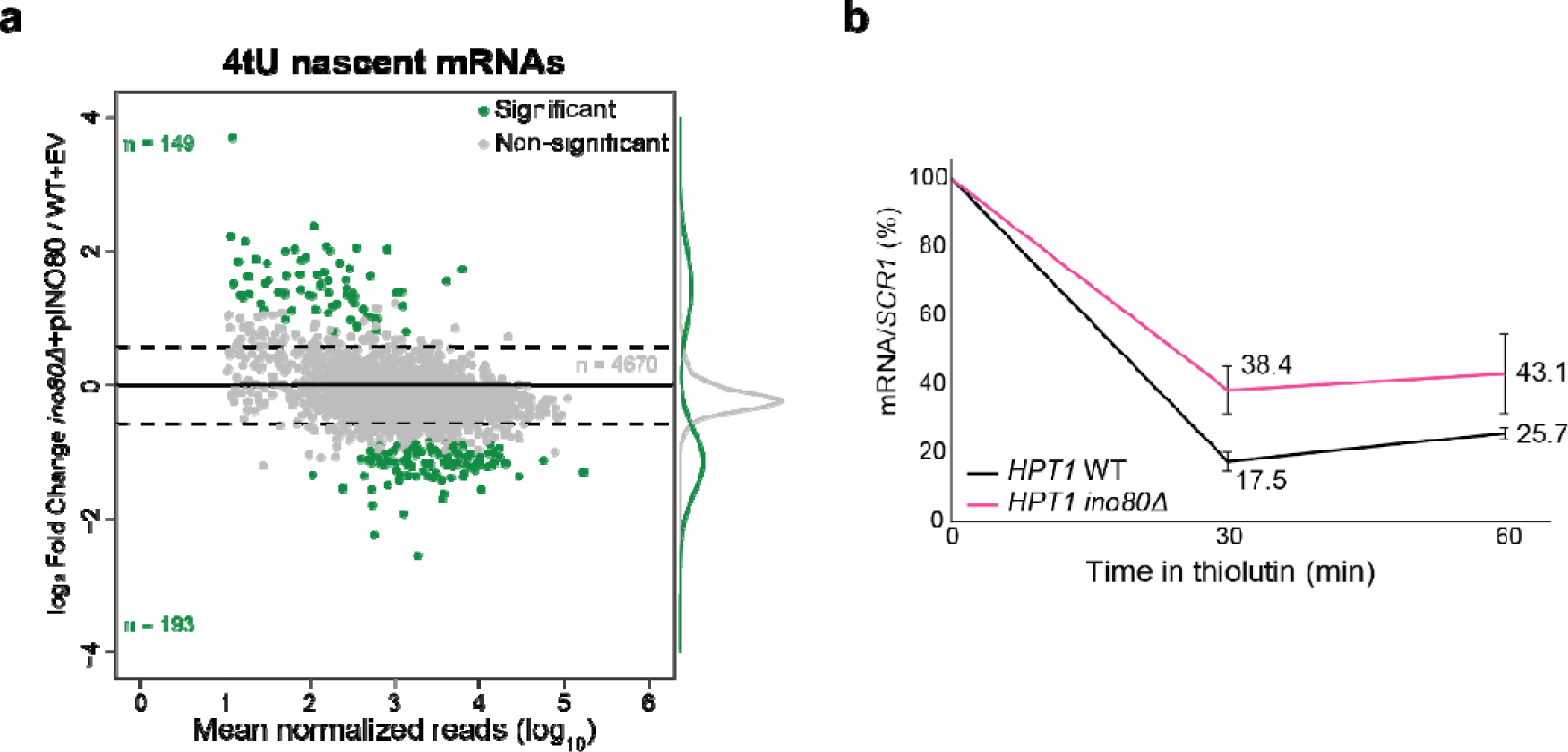
Ectopic expression of INO80 rescues the nascent mRNA synthesis defect of *ino80*Δ cells. **a,** Scatterplot analysis for fold changes in newly synthesized mRNA between *ino80*Δ cells expressing a WT allele of *INO80* from the pRS416 plasmid (*pINO80*) and WT cells transformed with the empty vector pRS416 (EV) plotted on a log_2_ scale. Analysis as in Fig.3a. **b,** Fold changes in mRNA decay rates between *ino80*Δ and WT for the indicated genes as calculated by cDTA analysis. **c,** RT-qPCR analysis for *HPT1* RNA was conducted in WT and *ino80*Δ before (T_0_) and after addition of thiolutin in YPD as in Fig. 1d.

**Extended Data Fig. 2.**
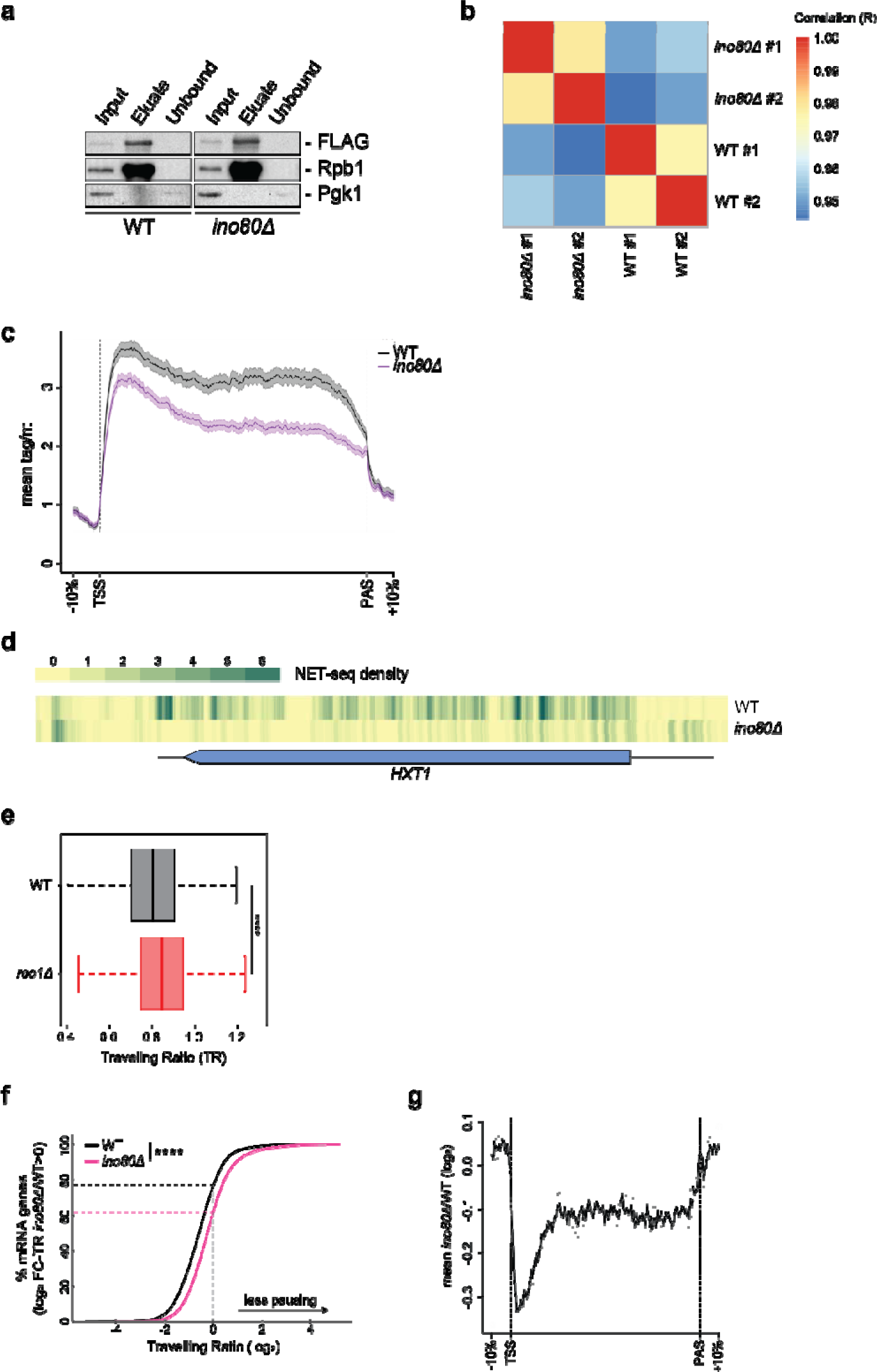
NET-seq analysis in the absence of INO80. **a,** Rpb1-FLAG is efficiently immunoprecipitated from yeast cells during NET-seq analysis. Lysates from WT and *ino80*Δ cells expressing *RPB1-FLAG* were subjected to FLAG immunoprecipitation using FLAG beads. Rpb1-FLAG was subsequently eluted from the beads with FLAG peptide. Input, eluate and unbound samples were immunoblotted against FLAG epitope and total Rpb1. Immunoblotting against Pgk1 serves as loading control. All samples were resolved in the same SDS-page gel. **b,** Heatmap of Pearson’s correlation coefficients (R) for all pairwise combinations of NET-seq reads in independent biological replicates. **c,** Metagene analysis for NET-seq read densities in WT and *ino80*Δ across mRNA genes from Fig. 2b. TSS: Transcription Start Site. TES: Transcription End Site. **d,** Genome browser snapshots of NET-seq read densities across the *HXT1* gene in WT and *ino80*Δ. **e,** Boxplot analysis comparing Traveling Ratios between WT and *rco1*Δ. **f,** Cumulative distribution plots comparing the Traveling Ratio (TR) values in WT and *ino80*Δ for mRNA genes with increased Traveling Ratio values in the absence of INO80 (log2 FC-TR *ino80*Δ*/*WT>0). ****p < 0.0001 with Mann-Whitney U test. **g,** Metagene analysis for the fold change in NET-seq read densities between *ino80*Δ and WT across mRNA genes with increased Traveling Ratios in the absence of INO80.

**Extended Data Fig. 3.**
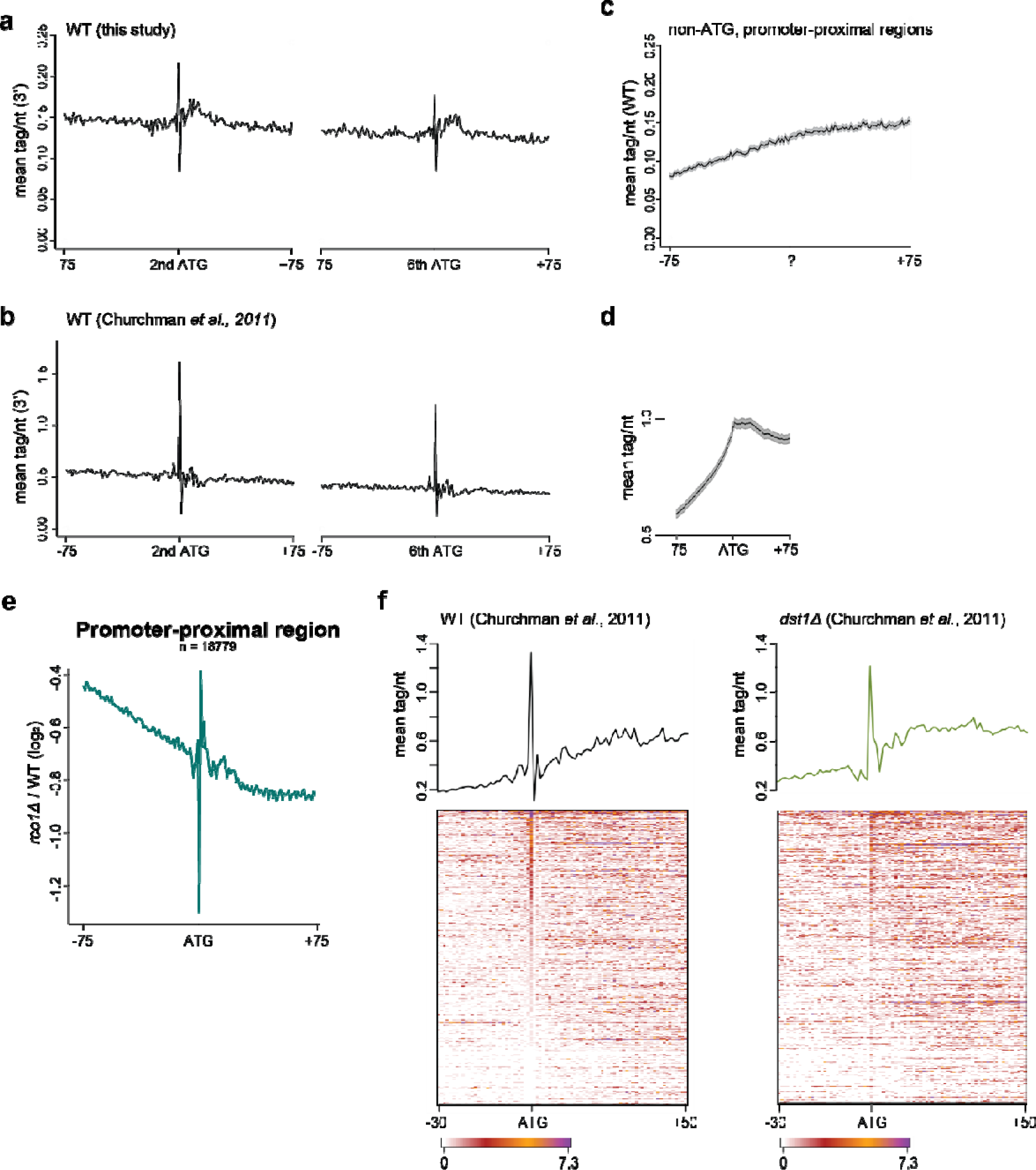
Pol II density at ATG sites. **a, b,** Metagene analysis for NET-seq read densities in WT cells from this study (a) and the Churchman et al., 2011 study^50^ (b) aligned at the A nucleotide of either the 2^nd^ or 6^th^ ATG motif found downstream the TSS. **c,** Metagene analysis for NET-seq read density in WT aligned at the first nucleotide of random trinucleotide sites in the promoter-proximal (TSS to TSS+200bp) regions. **d,** Metagene analysis for Pol II CRAC read densities aligned at the A nucleotide of the ATG sites in the promoter-proximal (TSS to TSS+200bp). **e,** Fold change metagene analysis comparing the NET-seq densities between WT and *rco1*Δ at the promoter-proximal ATG sites. **f,** Metagene and heatmap analysis for NET-seq read densities in WT (left) and *dst1*Δ (right) aligned at the A nucleotide of the 1^st^ ATG motif found downstream the TSS.

**Extended Data Fig. 4.**
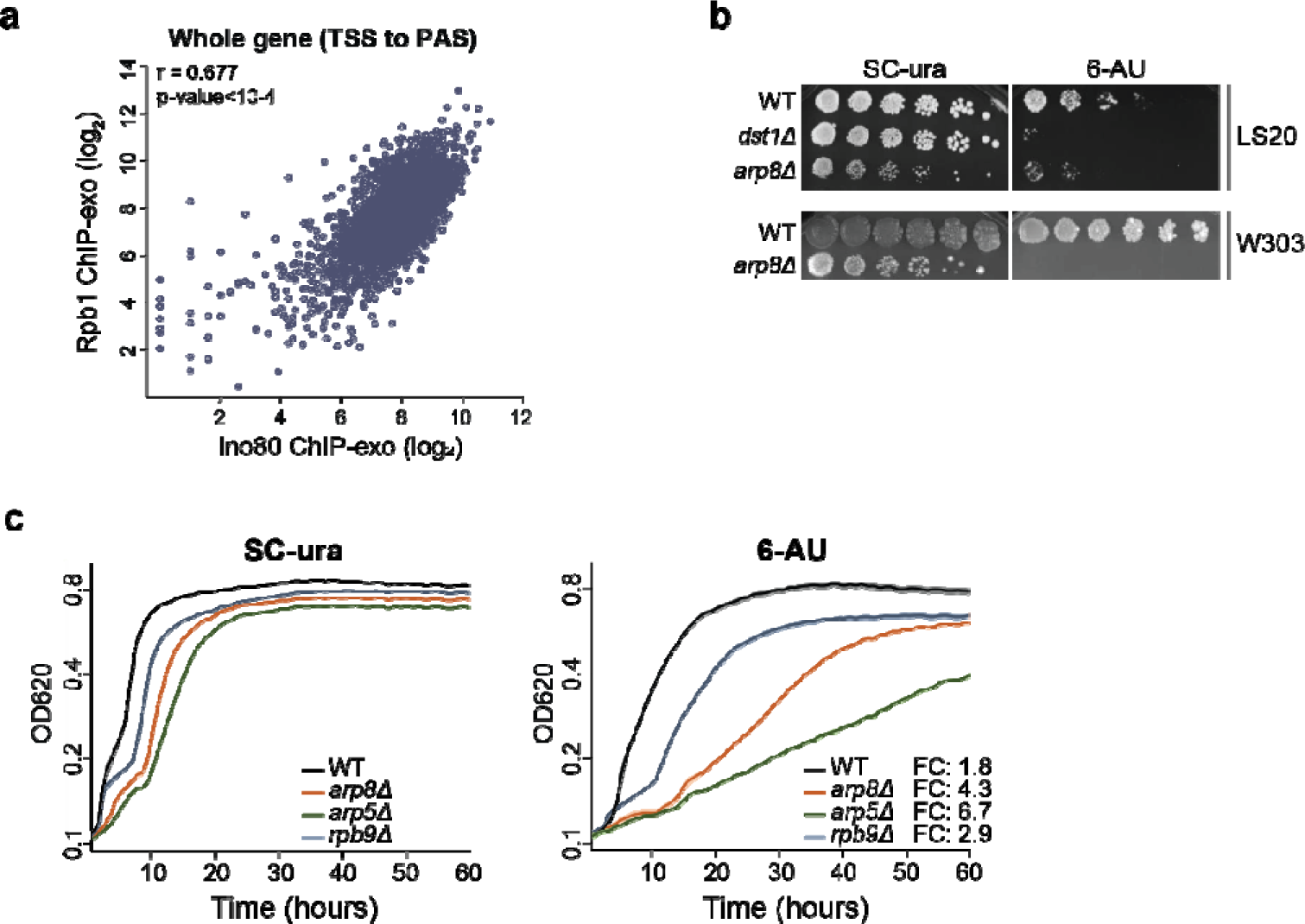
INO80 promotes the response to transcription elongation stress. **a,** Scatterplot analysis comparing Ino80^40^ and Rpb1 ChIP-exo read densities between the transcription start site and polyadenylation site (TSS to PAS) in the 5798 mRNA yeast genes. Pearson’s correlation coefficient (r) and p-value are indicated. **b,** 5-fold serial dilution of cells from the indicated strains were plated onto both SC-ura and SC-ura + 50µg/ml 6-Azauracil (6-AU) and incubated at 30°C for 4-5 days. Genetic backgrounds (LS20, W303) are indicated. The mutant strain deleted for the Dst1/TFIIS transcription elongation factor (*dst1*Δ) has been reported to have transcriptional elongation defects in 6-AU^87^ and was used as control. **c,** Cell proliferation analysis for the indicated strains grown exponentially in SC-ura (left panel) and in SC-ura containing 50µg/ml 6-azauracil (6-AU) (right panel) liquid media for the indicated time. Cell density was measured at OD^620^ and shown on a log_2_ scale. Shading surrounding each line denotes standard error from four independent experiments. The mutant strain deleted for the RNA Polymerase II subunit Rpb9 (*rpb9*Δ) has been reported to have transcriptional elongation defects in 6-AU^88^ and was used as control. Doubling time from OD^620^ 0.2 to OD^620^ 0.4 was calculated for all strains in both conditions. Fold Change ratio (FC) of doubling time in 6-AU over SC-ura is shown (right panel).

**Extended Data Fig. 5.**
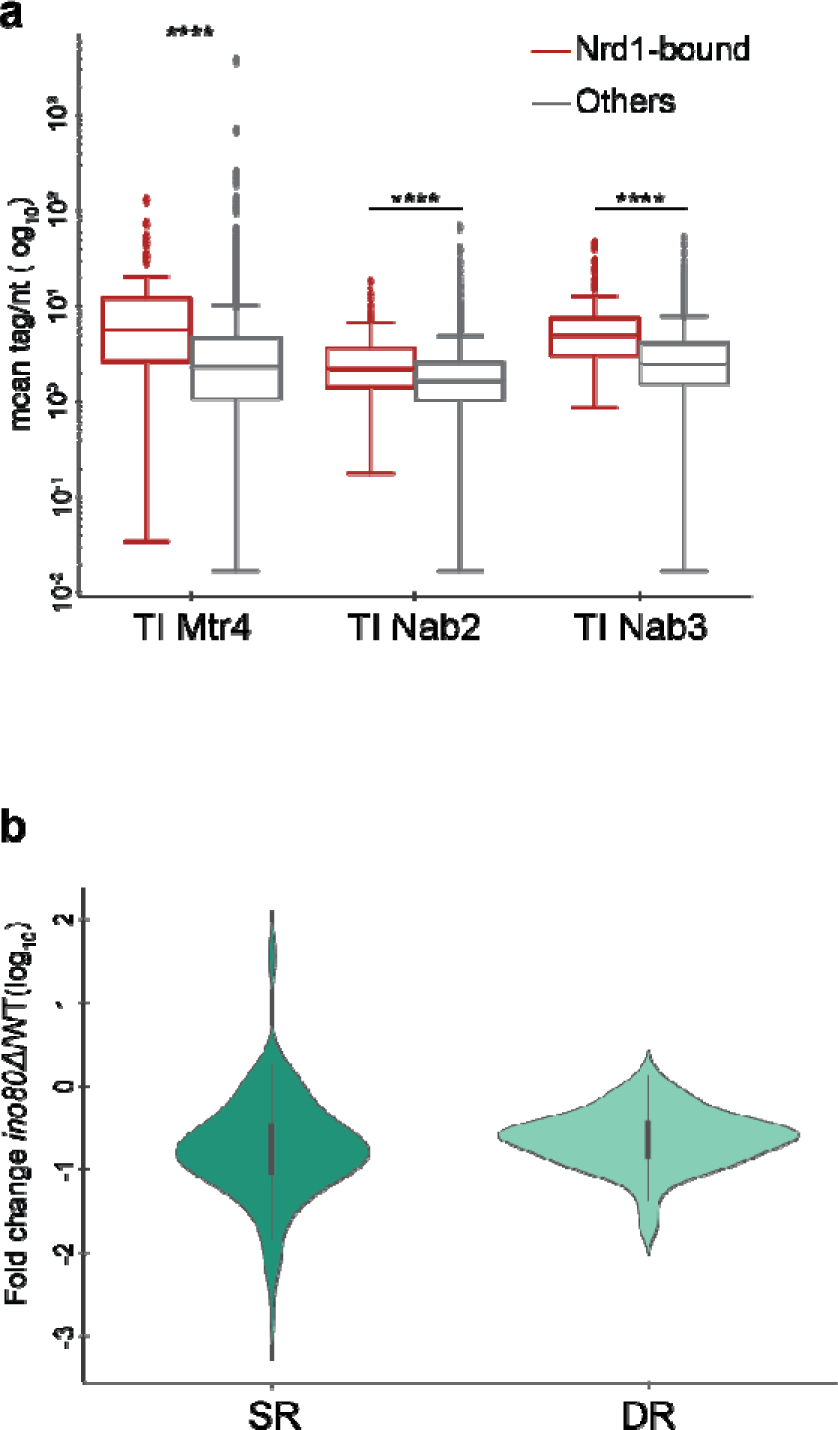
Analysis of Nrd1-bound genes. **a,** Boxplot analysis comparing Transcript Instability values for Mtr4, Nab3 and Nab2 calculated for the promoter-proximal regions in the Nrd1-bound and Others groups of genes. ****, p<0.0001 by Wilcoxon rank-sum test. **b,** Violin plot for the fold change in Synthesis rates (SR) and Decay rates (DR) between *ino80*Δ and WT, as calculated by cDTA analysis.

**Extended Data Fig. 6.**
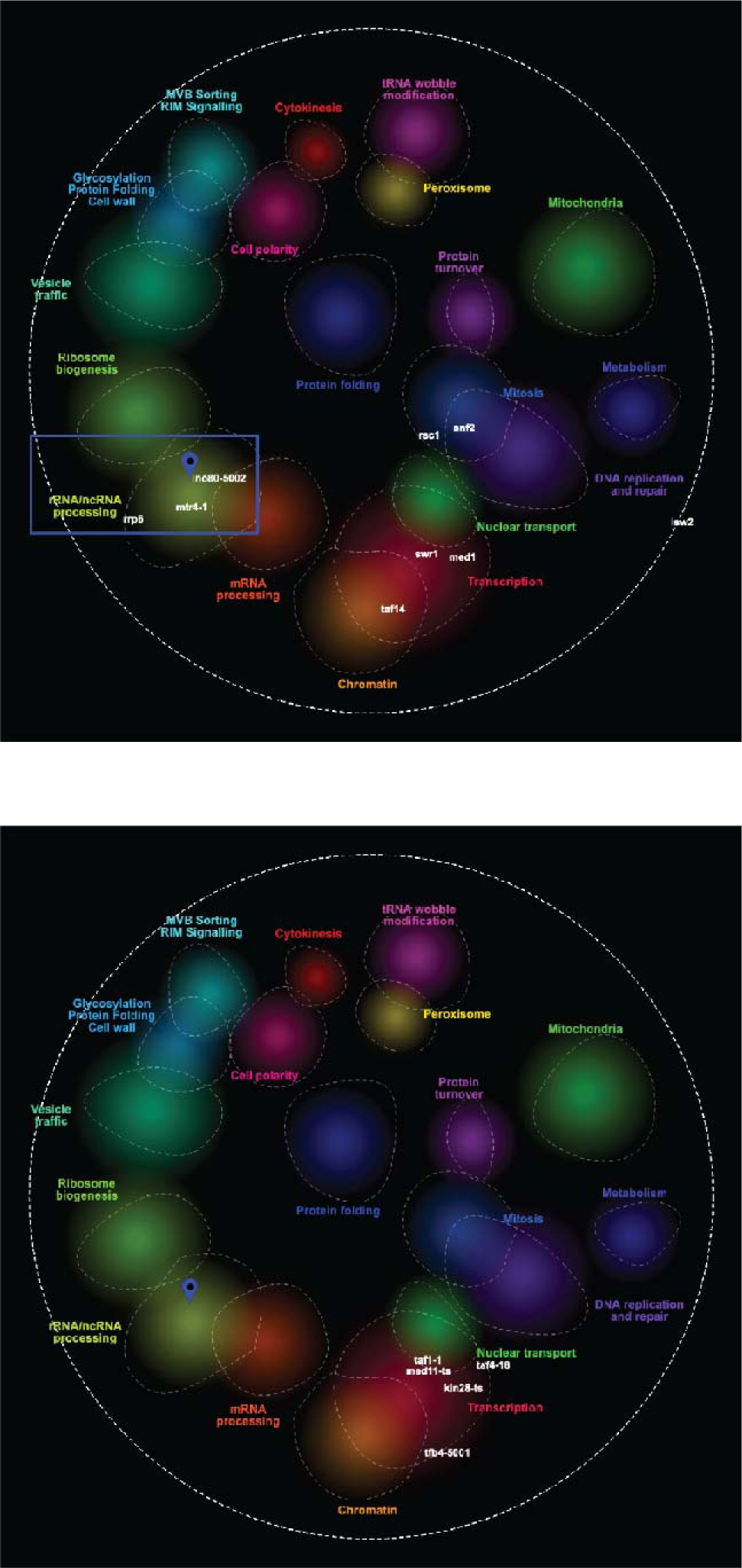
High throughput genetic interaction analysis of *INO80*. Top panel: Mapping of *INO80* on the CellMap global genetic interaction profile similarity network. The different functional cellular pathways are highlighted on the map. *INO80 (ino80-5002)* location is shown as a teardrop icon. The position of *INO80* within the non-coding RNA Processing pathway is highlighted in the blue rectangle. The positions of the non-coding RNA factors *RRP6* and *MTR4* (*mtr4-1*), the ATP-dependent chromatin remodelers *RSC1, SNF2, ISW2* and *SWR1,* and the transcription factors *MED1* and *TAF14* are shown on the map. Bottom panel: Mapping of the transcription factors *TAF1* (*taf1-1*), *MED11* (*med11-ts*), *KIN28* (*kin28-ts*)*, TFB4 (*tfb4*-5001), TAF4* (*taf4-18*) on the CellMap global genetic interaction profile similarity network.

**Extended Data Fig. 7.**
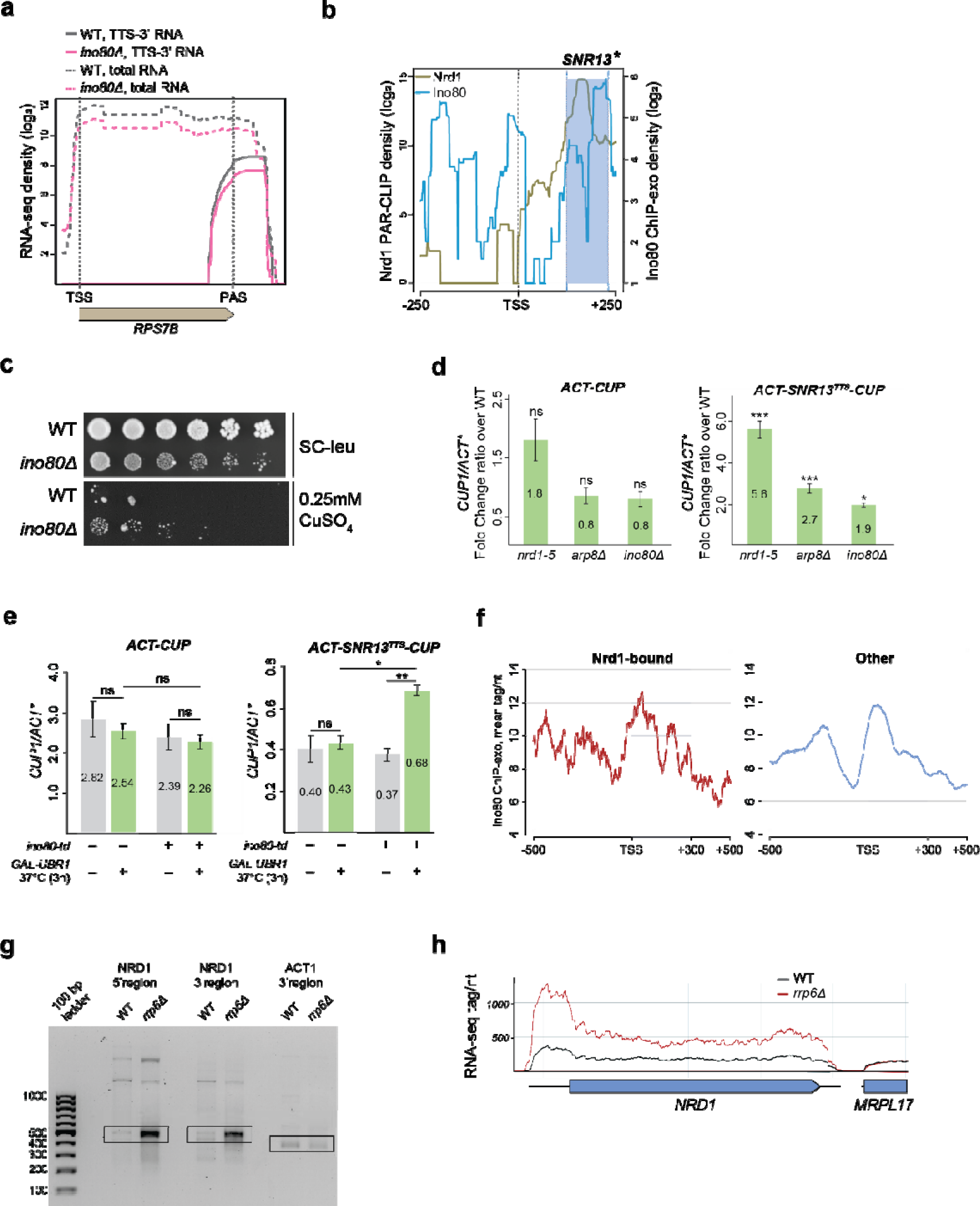
Defective NNS-dependent transcription termination in *ino80*Δ cells. **a,** Genome browser snapshot comparing RNA-seq read densities in WT and *ino80*Δ across the protein-coding gene *RPS7B*. Solid lines, density of reads encompassing the transcription termination site (TTS-3’). Analysis was performed around the polyadenylation site (PAS). Broken lines, total RNA-seq density. Genomic coordinates of Chromosome XVI around the *RPS7B* gene are shown. The closest gene downstream of *RPS7B* is the *PHO23* mRNA gene (TSS: Chromosome XIV, 442509). **b,** Genome browser snapshot of Nrd1 PAR-CLIP density profile from ^89^ and Ino80 ChIP-exo density profile from ^40^ across the snoRNA *SNR13* gene. Tags were aligned relative to TSS and plotted on a log_2_ scale. The region between +125 to +232bp from TSS cloned in the *ACT-SNR13^TTS^-CUP* plasmid^8^ is highlighted in blue (*SNR13\**). **c,** 5-fold serial dilution of cells from the indicated strains were plated onto both SC-leu and SC-leu + 0.25mM CuSO_4_ and incubated at 30°C for 3 (SC-leu) and 7 days (CuSO_4_). **d,** RT-qPCR analysis for the *CUP1*/*ACT\** ratio was conducted in the indicated strains transformed with either the *ACT-CUP* (left panel) or *ACT-SNR^TTS^-CUP* (right panel) plasmids. Bars, standard errors from three independent experiments. Asterisks indicate statistical significance of the change in the respective mutant compared to the WT as calculated by t-test. ns: non-significant. *, p<0.05. ***, p<0.001. **e,** RT-qPCR analysis was conducted in both WT and the *ino80-td* degron-inducible strain (*ino80-td*) ^68^ transformed with the *ACT-CUP* control plasmid described in a. Galactose-induced activation for 3 hours at 37°C (+) and glucose-induced suppression (–) of the *UBR1* N-recognin in the two strains is indicated. Bars, standard errors from four independent experiments. p-values were calculated by t-test after testing for normal distribution by Shapiro-Wilk test. ns, non-significant. **f,** Metagene analysis for Ino80 ChIP-exo density from ^40^ across the Nrd1-bound (left panel) and Others (right panel) groups of genes. Tags were aligned to the Transcription Start Site (TSS) for each gene. The promoter-proximal region enriched in Ino80 in the Nrd1-bound genes is highlighted in blue **g,** Representative image of nested 3’RACE analysis of termination events at the *NRD1* 5’ and 3’ regions according to the scheme in Figure 7g and at the 3’ of *ACT1* gene in WT and *rrp6*Δ on an agarose gel. Boxes indicate the most frequent termination sites that were analysed by ImageJ. The 100bp ladder is shown. *ACT1* serves as loading control. **e,** Genome browser snapshot of RNA-seq read distribution across the *NRD1* gene in WT and *rrp6*Δ.

**Extended Data Fig 8.**
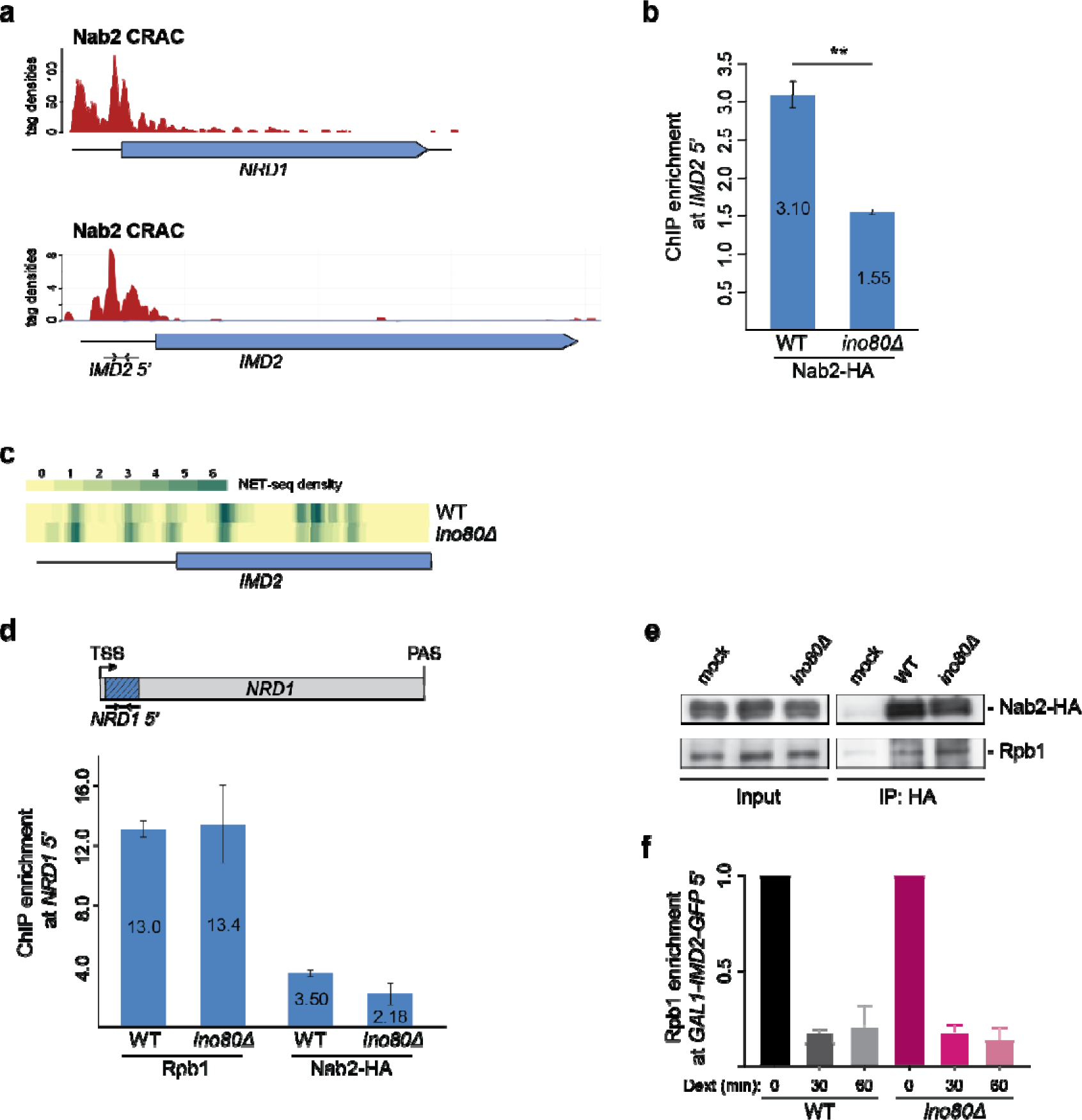
Co-transcriptional recruitment of Nab2 to chromatin and release of Pol II for non-coding termination depend on INO80. **a,** Genome browser snapshot for CRAC-seq read densities for Nab2 from ^4^ across the *NRD1* (top) and the *IMD2* (bottom) genes. **b,** Chromatin immunoprecipitation for Nab2-HA in WT and *ino80*Δ cells. Fold enrichment analysis at the promoter-proximal region of the *IMD2* gene (*IMD2 5’*), as depicted in (a). **c,** Genome browser snapshot for NET-seq read densities at the promoter-proximal region of the *IMD2* gene in WT and *ino80*Δ. **d,** Chromatin immunoprecipitation for Rpb1 and Nab2-HA in WT and *ino80*Δ cells. Fold enrichment analysis at the promoter-proximal region of the *NRD1* gene (*NRD1 5’*) (upper panel, shaded in dark blue) was conducted as in Fig. 8a. **e,** Nucleic acid-free lysates from WT and *ino80*Δ cells both expressing Nab2-HA were subjected to HA-IP. Inputs and IP samples were immunoblotted for Rpb1 and HA. Mock, WT protein lysate subjected to immunoprecipitation with a control IgG antibody. **f,** Chromatin immunoprecipitation for Rpb1 in WT and *ino80*Δ cells grown in galactose and after 30 and 60 min of dextrose addition in the galactose-containing medium. Fold enrichment analysis at the promoter-proximal region of the *GAL1-IMD2 TE-GFP* gene (*5’*) was conducted as in Fig. 8e.

**Extended Data Table 1.**
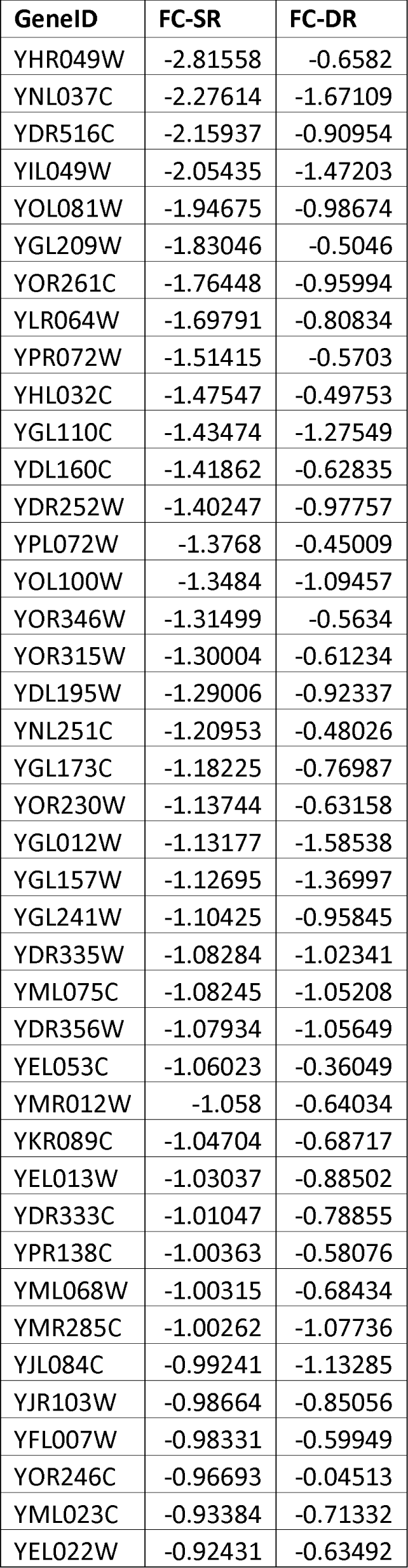
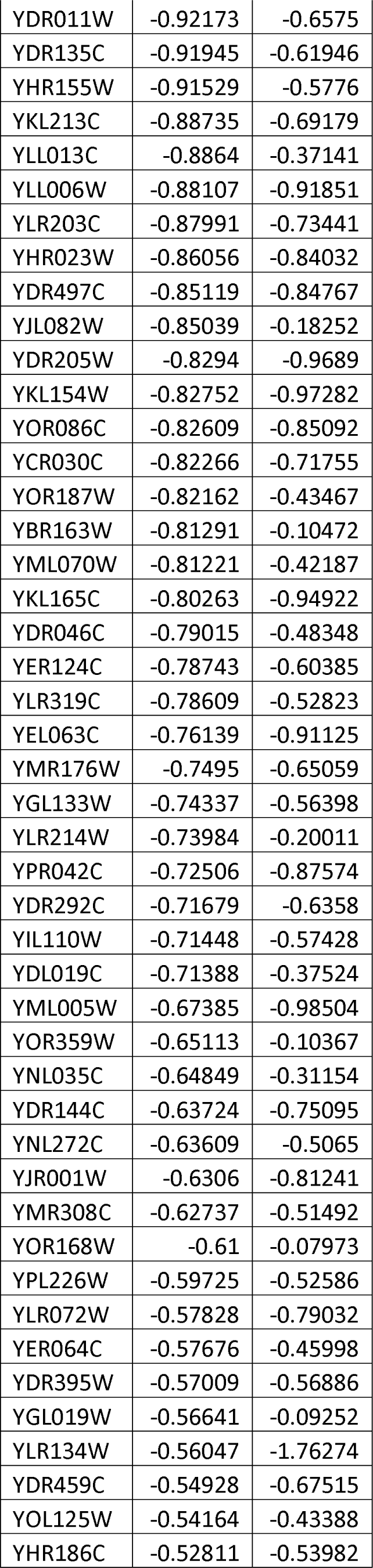
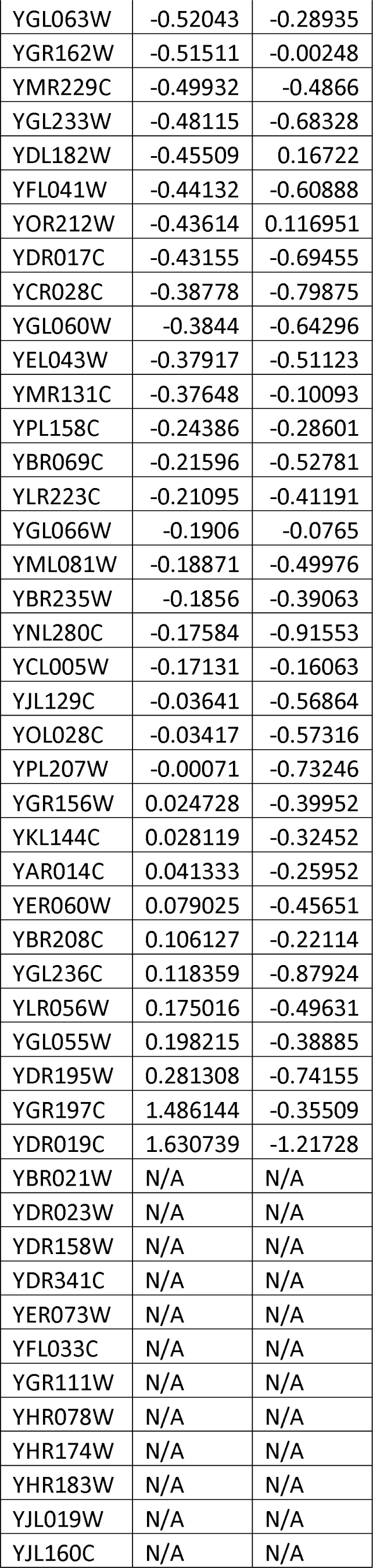
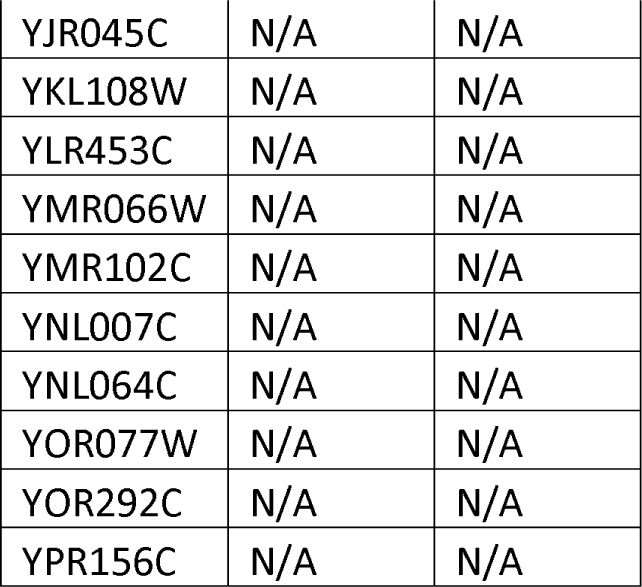
List of Nrd1-bound genes identified in ^27^ with log2 fold change synthesis and decay rates (FC-SR and FC-DR respectively) in *ino80*Δ compared to WT.

**Extended Data Table 2.** Expression of genes encoding for RNA surveillance proteins is similar in WT, *ino80*Δ and *arp8*Δ cells. RNA abundance for the listed genes was measured by spiked-in total RNA-seq analysis in the indicated strains. Values represent average from two biological replicates for each experiment. The RNA-seq experiment for WT(1) and *ino80*Δ was conducted independently from WT(2) and arp8Δ. Values represent average from two biological replicates for each experiment.

| Pathway/complex | Gene | WT<sup>(1)</sup> | *ino80Δ* | Fold Change | WT<sup>(2)</sup> | *arp8Δ* | Fold Change |
| --- | --- | --- | --- | --- | --- | --- | --- |
| **NNS** | *NRD1* | 0.61 | 0.67 | **1.1** | 0.43 | 0.37 | **0.8** |
|  | *NAB3* | 0.54 | 0.44 | **0.8** | 0.30 | 0.30 | **1.0** |
|  | *SEN1* | 0.46 | 0.58 | **1.3** | 0.29 | 0.25 | **0.8** |
| **Exosome** | *RRP6* | 0.68 | 0.54 | **0.8** | 0.46 | 0.46 | **1.0** |
|  | *DIS3/RRP44* | 0.82 | 0.73 | **0.9** | 0.64 | 0.62 | **1.0** |
| **TRAMP** | *MTR4* | 0.97 | 0.74 | **0.8** | 0.87 | 0.89 | **1.0** |
|  | *TRF4* | 0.45 | 0.48 | **1.1** | 0.28 | 0.32 | **1.1** |
|  | *AIR1* | 0.38 | 0.30 | **0.8** | 0.29 | 0.29 | **1.0** |
|  | *AIR2* | 0.21 | 0.22 | **1.1** | 0.11 | 0.15 | **1.4** |
| **Surveillance, cleavage and polyadenylation** | *HRP1* | 1.21 | 1.01 | **0.8** | 1.04 | 0.83 | **0.8** |
|  | *PCF11* | 0.26 | 0.24 | **0.9** | 0.19 | 0.18 | **1.0** |
|  | *NAB2* | 0.70 | 0.59 | **0.8** | 0.51 | 0.54 | **1.1** |
|  | *PAB1* | 8.95 | 6.85 | **0.8** | 4.53 | 3.89 | **0.9** |
| **Capping** | *CEG1* | 0.42 | 0.44 | **1.1** | 0.19 | 0.22 | **1.1** |
|  | *CET1* | 0.27 | 0.26 | **1.0** | 0.17 | 0.19 | **1.1** |
| **Early mRNA processing** | *CBC1* | 0.96 | 0.74 | **0.8** | 0.58 | 0.68 | **1.2** |
|  | *GBP2* | 1.43 | 1.09 | **0.8** | 0.83 | 0.74 | **0.9** |
|  | *THO2* | 0.48 | 0.51 | **1.0** | 0.27 | 0.31 | **1.2** |
| **Export** | *MEX67* | 0.52 | 0.50 | **1.0** | 0.32 | 0.34 | **1.1** |
| **Cytoplasmic decay** | *XRN1* | 1.87 | 1.76 | **0.9** | 0.91 | 0.91 | **1.0** |
|  | *SKI2* | 0.40 | 0.40 | **1.0** | 0.28 | 0.31 | **1.1** |

**Extended Data Table 3.** Yeast strains used in this study.

| Name | Genotype | Source |
| --- | --- | --- |
| YM50 | LS20 [*MatΔ ade2 can1r cyh2r lys S URA3-52 trp1Δ his3-Δ200GAL:HO leu 2Δ*] | Sandell & Zakian, *Cell* 1993 |
| YM85 | LS20 *ino80Δ::KANMX* | Papamichos-Chronakis & Peterson, *Nat Struct Mol Biol* 2008 |
| YM133 | W303 *ubr1Δ::HIS3-GAL-Myc-UBR1* | Jónsson *et al.*, *Mol Cell* 2004 |
| YM135 | W303 *ubr1Δ::HIS3-GAL-Myc-UBR1 ino80Δ::URA3-HA-ino80-td* | Jónsson *et al.*, *Mol Cell* 2004 |
| YM181 | LS20 *arp8Δ::KANMX* | Papamichos-Chronakis *et al.*, *Cell* 2011 |
| YM311 | W303 [*MATa ade2-1 can1-100 his3-11 leu2,3,112 trp1-1 ura3-1*] | Verma *et al.*, *Mol Cell* 2011 |
| YM334 | W303 *arp8Δ::KANMX* | Lafon *et al.*, *Mol Cell* 2015 |
| YM467 | LS20 *dst1Δ::KANMX* | This study |
| YM468 | LS20 *rpb9Δ::KANMX* | This study |
| YM753 | BY4741 [*MATa his3Δ1 leu2Δ0 met15Δ0 ura3Δ0*] *NRD1-3XHA::HIS3* | This study |
| YM801 | 46a [*MATa ura3 his3 trp1 lys2 ade2 leu2 cup1Δ*] | Steinmetz & Brow, *Mol Cell Biol* 1996 |
| YM802 | 46a *nrd1-5* | Steinmetz & Brow, *Mol Cell Biol* 1996 |
| YM838 | LS20 *arp5Δ::KANMX* | This study |
| YM850 | BY4741 *PCF11-3XHA::HIS3* | This study |
| YM885 | LS20 *RPB1-3XFLAG::KANMX* | This study |
| YM900 | BY4741 *PCF11-3XHA::HIS3 INO80-5XFLAG::NATMX* | This study |
| YM901 | BY4741 *NRD1-3XHA::HIS3 INO80-5XFLAG::NATMX* | This study |
| YM903 | LS20 *RPB1-3XFLAG::KANMX ino80Δ::HPHMX* | This study |
| YM1002 | 46a *arp8Δ::HPHMX* | This study |
| YM1036 | 46a *nrd1-5 arp8Δ::HPHMX* | This study |
| YM1139 | LS20 *rrp6*Δ::*KANMX* | This study |
| YM1325 | LS20 *NAB2-3XHA*::*KANMX* | This study |
| YM1331 | LS20 *NAB2-3XHA*::*KANMX* *ino80*Δ::*HPHMX* | This study |
| YM1332 | LS20 *NAB2-3XHA*::*KANMX* *ino80*Δ::*HPHMX* | This study |

## METHODS

### Bioinformatics Tools and genome annotations

Unless otherwise indicated, version of the tools used for all bioinformatics analyses are as follows:

Samtools version 1.10

Bedtools version 2.29.2

Bowtie2 version 2.4.1

Hisat2 version 2.2.0

R version 3.6.2

BamCoverage version 3.5.0

Fastqc version 0.11.9

Cutadapt version 2.10

Trim_galore 0.6.4

MarkDuplicates version 2.23.1

FeatureCounts version 2.0.0

The *Saccharomyces cerevisiae* reference genome (S288C version R64-2-1) was downloaded from Saccharomyces Genome Database (https://www.yeastgenome.org/). The annotation used in the analysis for mRNA genes was taken from ^90^XUT annotation was produced by the Morillon lab^91^. SUT and CUT annotations come from Xut et al (2009) and NUT genes from ^22^. Annotations for other transcript types (snRNA, snoRNA, etc) were downloaded from Saccharomyces Genome Database.

The *Schizosaccharomyces pombe* reference genome and annotation (ASM294v2.47) were downloaded from pombase (www.pombase.org). Only protein coding genes were used in this analysis.

### ATG analysis

ATG positions were found in S. cerevisiae genome using “grep” command, with pattern ATG or its reverse complement. These positions were then intersected with mRNA TSS region (TSS to TSS+200) or gene body region (TSS+200 to TTS) using bedtools.

### NET-seq analyses

NET-seq was performed in duplicate from yeast strains containing Rpb1-FLAG (3XFLAG), alone or along with deletion of *INO80*. All strains appeared fully functional, with identical Ser2 and Ser5 phosphorylation levels on Rpb1 C-terminal domain and identical growth rates to the respective untagged strains (data not shown). NET-Seq libraries were constructed from biological duplicates of YM903 (*ino80*Δ, *RPB1-FLAG*) and YM885 (WT, *RPB1-FLAG*) cells and sequenced as previously described^92^. Reads shorter than 10 nt were filtered out. Reads were mapped with Bowtie2 using default parameters. Duplicates reads were marked using Picard MarkDuplicates. Coverage files (bigWig) were produced by custom script. Reads locations from the alignment files (bam) were turned into bed files using bedtools bamtobed (with « -split » option), ignoring duplicates reads and reads with mapping quality below 30, piled up along the genome and turned into bigWig files using a python script. FeatureCounts was used for quantification with options « -O --fraction s 2 --ignoreDup ». Normalization factors were computed using DESeq2 method, using the whole count matrix. Travelling Ratio was computed as: (read density on gene body) / (read density on TSS), with gene body the region between TSS+200 nt to TTS and TSS the region from TSS to TSS + 200 nt. Read density was computed as: (# reads on region / region length) *1000.

For the ATG analysis, only the position of the first nucleotide at the 5’ end of the mapped read was used, which correspond to the 3’ end of the sequenced fragment.

### 4tU labeling and purification of newly synthesized RNAs

RNA labeling and purification were performed in triplicate as previously described with some minor adjustments^46,93^. Briefly, 20 mL of wildtype and mutant S. cerevisiae cells were grown at 30°C to an OD600 ≈0.8 in YPD. Newly synthesized (ns) RNAs were labelled for 6 min with 4-thiouracil (4tU, Sigma-Aldrich) added to the yeast cultures to a final concentration of 5 mM. Similarly, S. pombe RNAs were labeled for 6 min in YES medium at 32°C. Cells were pelleted directly after labeling and flash frozen in liquid nitrogen. All experiments were performed in biological triplicates. Prior to RNA extraction S. cerevisiae and S. pombe cells were mixed in a ratio of 3:1 and total RNA was extracted with Ribopure Yeast Kit (Ambion, Life Technologies) according to the manufacturer’s guidelines. 150 µg of total RNA were biotinylated for 3 hours at room temperature using 200 μL of 1 mg/mL EZ-link HPDP-Biotin (Pierce) with 100 μL of biotinylation buffer (100 mM Tris-HCl pH 7.5, 10 mM EDTA) and 100 μL of DMSO adjusted to a final volume of 1 mL with DEPC-treated RNase-free water (Sigma-Aldrich). After biotinylation, excess biotin was removed by mixing 1 mL of chloroform to the reaction and the liquid phases were separated by centrifugation for 5 min at 4°C and max speed. The aqueous phase was recovered and precipitated with isopropanol (1/10 volumes 5M NaCl and 2.5 volumes isopropanol). The recovered RNA was resuspended in 100 µL DEPC-treated RNase-free water and incubated for 10 min at 65°C followed by 5 min on ice. nsRNAs were bound to 100 μL of μMACS streptavidin microbeads (Miltenyi Biotec) for 90 min at room temperature and purification was performed using the μMACS streptavidin starting kit (Miltenyi Biotec). Columns were first equilibrated with 1 mL of washing buffer (100 mM Tris-HCl at pH 7.5, 10 mM EDTA, 1M NaCl, 0.1% Tween20). Samples were loaded and the flow-through was reapplied 2 times to the columns. The columns were washed five times with increasing volumes of washing buffer (600, 700, 800, 900, and 1000 μL). Ultimately, labeled RNAs were eluted twice with 200 μL of 100mM DTT. nsRNAs were precipitated overnight in 1/10 volume of 3 M NaOAc, 3 volumes of 100% ethanol and 20 μg of RNA-grade glycogen. The RNAs were recovered by centrifugation and RNA pellets were wash with ice-cold 70% ethanol and resuspended in 20 μL of DEPC-treated RNase-free water (Sigma-Aldrich). Samples were stored at −80°C until further use. Mapping of the reads was performed with Hisat2 on a merged reference genome of S. cerevisiae and S. pombe with options “-5 10 --maxintronlen 5000”, and reads mapping on both genomes and/or with mapping quality < 30 were removed. FeatureCounts was used for quantification, with options “-O –C –M -Q 30 --fraction --primary -s 2 -p”. Coverage was computed using bamCoverage (deepTools) with options “--binSize 1 --skipNAs” for reads with mapping quality > 30. Differential gene expression analysis was performed using DESeq2 1.16.1 Bioconductor R package on Saccharomyces cerevisiae counts normalized with size factors computed by the median-of-ratios method proposed by Anders and Huber on *Schizosaccharomyces pombe* counts (Anders and Huber 2010). cDTA analysis was performed as described^94^.

### Total RNA-seq

Yeast strains were processed in duplicate. ERCC RNA Spike-in control mixes (Ambion, catalogue #4456740) were added to 1µg of total RNA according to manufacturer. Ribosomal RNAs were depleted from total RNA using the RiboMinus™ Eukaryote v2 Kit (Life Technologies). Depletion efficiency and quality control of rRNA-depleted RNA was assessed by analysis in RNA Pico 6000 chip for 2100 Bioanalyzer (Agilent). Total RNA-seq libraries were prepared from 50ng of rRNA-depleted RNA using the TruSeq® Stranded Total RNA Sample Preparation Kit (Illumina). Paired-end sequencing (2×50 nt) was performed on a HiSeq 2500 sequencer giving between 65 and 84 million reads. Reads were mapped using version 2.0.6 of TopHat, with a tolerance of 3 mismatches and a maximum size for introns 2Kb. Tags densities were normalized on spike-in controls for all subsequent analyses.

### ChIP-exo

ChIP-exo was performed according to ^59^. In brief, yeast strains were grown to exponential phase in yeast extract peptone (YP) + 2% dextrose (30°C to OD600 nm = 0.8), then subjected to 1% formaldehyde crosslinking for 15 min at 25 °C. After quenched with 125mM final concentration of glycine for 5 min, cells were harvested and washed. Sonicated chromatin was prepared by standard methods. Standard ChIP methods were used for Rpb1 (y-80 Santa Cruz) followed by lambda exonuclease treatment and library construction. Libraries were sequenced on an Illumina 2000 sequencer. Reads were uniquely mapped to the reference genome (sacCer3 build) using version 2.1.0 of Bowtie2 with a tolerance of 1 mismatch in seed alignment. Data were normalized on the total number of uniquely mapped reads, for each sample.

### Cell proliferation analysis in liquid media

Liquid growth rate analysis was conducted using the CG-12 robot (S&P Robotics Inc.). Yeast cells transformed with a URA3-containing plasmid (pRS416) and grown exponentially in SC-ura at 30°C were diluted to OD620 0.1 in either SC-ura or SC-ura + 50µg/ml 6-AU and four 150µl samples from each strain were plated onto 96-well transparent plates covered with breathable films. Plates were incubated at 30°C for several days and OD620 was measured in the CG-12 robot (S&P Robotics Inc.) every hour with a shaking of 1000rpm for 1 minute before each read. Liquid growth rate analysis was not possible in the *nrd1-5* and associated strains due to flocculation. All cell proliferation experiments were repeated at least twice.

### Growth rates analysis on agar plates

Growth of yeast strains was compared by plating 5-fold serial dilutions on SC / SC-inositol, SC-ura / SC-ura + 50µg/ml 6-azauracil (6-AU, Sigma), SC-leu / SC-leu + 0.4mM CuSO4 (Sigma) or SC-leu / SC-leu + 0.25 CuSO4 and incubating the plates at 30°C for several days. All plating experiments were performed at least three times.

### RT-qPCR

Total RNA was extracted either with RNeasy Mini Kit (Qiagen) or by classical acid phenol-chloroform extraction. RNAs abundance was measured with SuperScript® III Platinum® SYBR® Green One-Step qRT-PCR Kit (Invitrogen) using specific primers for the targets of interest in a 10µl reaction volume, on Applied Biosystem Step-One Plus machines. RNAs abundance of each sample was compared with standard curves analysis. Abundance of *CUP1* from transcriptional read-through analyses using the *ACT-CUP* reporter system was compared to *ACT\** RNA specifically transcribed from the same plasmid. Abundance of *NRD1* promoter-proximal RNA (*NRD1* 5’) was compared to levels of *NRD1* promoter-distal RNA (*NRD1* 3’).

### 3’ RACE

The 3’ RACE kit from Invitrogen (18373-019) was used according to the instructions. Briefly, DNA-free acid-phenol extracted RNA was purified from exponentially grown yeast cells and reversed transcribed with Superscript II using the 3’ RACE Adaptor Primer oligo. RNA was subsequently degraded, and exponential PCR amplification was performed on the cDNA using a nested, gene specific primer and the AUAP primer from the kit. Products were run on a 1.5% agarose gel and visualized by Gelite™ X100 DNA Gel Stain from Stratech (17701-AAT). Quantitation analysis was conducted with ImageJ. The optimal cycle number for PCR amplification of cDNA for each primer pair was selected by confirming the linear amplification of a 4-sample serial dilutions series of cDNA template from WT, covering a 10-fold concentration range. For both *NRD1* 5’ and *NRD1* 3’, PCR amplifications of the cDNA template were conducted in 23 cycles.

### Co-immunoprecipitation

Benzonase-treated yeast cell extracts were incubated with 20µl Anti-FLAG Affinity Gel (Sigma Aldrich) in 10mM Tris-Cl pH 7.5, 150mM NaCl, 0.05mM EDTA, 0.5% NP40, 1X Protease Inhibitor Cocktail (Roche) and 0.01mM PMSF for 3h. Beads were washed twice in 10mM Tris-Cl pH 7.5, 150mM NaCl, 0.05mM EDTA, 1X Protease Inhibitor Cocktail (Roche) and 0.01mM PMSF and resuspended in SDS sample buffer. Western blot analyses were performed using Monoclonal anti-FLAG M2 antibody (Sigma F3165), Anti-HA tag antibody (Abcam, ab9110) and Anti-Arp5 antibody (Abcam, 12099). Images were acquired using Syngene G:BOX. The experiment was reproduced three times. Immunoprecipitations for HA-tag were performed similarly on benzonase-treated yeast cell extract using 36µl Protein-A sepharose beads coupled with 6µl of appropriate antibody (Anti-HA tag, Abcam ab9110). Control immunoprecipitations (mock) were performed with a non-specific antibody (Anti-Rabbit IgG, Promega). Western blot analyses were performed using the following antibodies: Anti-HA, Anti-Arp5, and Anti-Rpb1 y-80 (Santa Cruz Biotechnology, sc-25758).

### Ino80 inducible degradation

The assay for inducible degradation of Ino80^68^ was performed as previously described^43,95^ Briefly, a degron tag was fused to INO80 (Ino80-td) in cells carrying the *UBR1* gene, that encodes for the N-recognin Ubr1, under the control of the *GAL1* promoter, and cells were transformed with the *ACT-CUP* plasmid reporter system. Cells were grown in SC-leu media containing raffinose as carbon source at 25°C until OD600∼0.6-0.7, then shifted to SC-leu media containing galactose to induce *UBR1* expression. Subsequently, complete Ino80-td protein degradation by the Ubr1 pathway was induced by 3 hours of heat shock at 37°C. Cells were then immediately harvested by centrifugation at 4°C, 3500rpm for 3 minutes, then washed twice with ice-cold water and snap-frozen in liquid nitrogen for RNA analysis. Yeast cells carrying the GAL1-driven UBR1 gene along with an unmodified INO80 genes were use as control. Furthermore, both (*GAL1-UBR1*) and (*GAL1-UBR1 INO80-td*) cells were grown in SC-leu media containing glucose as a negative control for UBR1 expression. RNA analysis was conducted by RT-qPCR as described above.

### ChIP-qPCR

Chromatin immunoprecipitation was performed as previously described in ^96^. Sonication was performed on a Bioruptor® Pico sonication device (Diagenode) with ten 30-sec pulses with a 30-sec break between pulses. Immunoprecipitation was carried out with Anti-HA tag antibody (Abcam ab9110) and Anti-Rpb1 y-80 antibody (Santa Cruz Biotechnology, sc-25758) in parallel. The recovered DNA was subjected to quantitative real-time PCR using Platinum® SYBR® Green qPCR SuperMix-UDG Kit (Invitrogen) with specific primers for the targets of interest in 10µl reaction volume, on Applied Biosystem Step-One Plus machines.

### RNA-IP

RNA immunoprecipitation was performed as described in ^97^. Sonication was performed on a Bioruptor® Plus sonication device (Diagenode) with two 15-sec pulses at 50% amplitude with a 1-min break between pulses. Immunoprecipitation was carried out with Anti-HA tag antibody (Abcam ab9110). The recovered RNA was analysed by RT-qPCR as described above.

### De novo analysis of public ChIP-exo and CRAC data

Nab3 and Pol II CRAC data were taken from GEO dataset GSE70191 and processed as described in ^57^. Nab2 and Mtr4 CRAC data were taken from GEO dataset GSE46742 and processed as described in ^4^. ChIP-exo Rpb1 data were downloaded and processed as described in ^41^.

